# Ultrastructurally planar growth cones during neocortical axon navigation

**DOI:** 10.64898/2026.10.05.756131

**Authors:** Yu Nakanishi, Masafumi Tsuboi, Hiroki Kawai, Koyo Shibayama, Kenta Yamauchi, Sasikarn Looprasertkul, Kyungeun Lee, Hajime Komaki, Takamitsu Egashira, Shogo Suga, Tomonari Murakami, Kenichi Ohki, Hiroyuki Hioki, Nobuhiko Ohno, Hidenobu Mizuno, Yusuke Hirabayashi

## Abstract

Neurons extend axons toward spatially defined but often distant targets to establish brain circuits. Callosal projection neurons exemplify this organization, linking corresponding cortical territories between the two hemispheres^1^. Growing axons rely on growth cones to regulate direction and speed through densely packed tissues^2,3^. Although growth cone morphology and dynamics have been extensively characterized in cultured neurons and invertebrate models, how growth cones navigate the three-dimensional environment of the developing mammalian brain remains poorly explored. Here, we combined a genetically encoded electron microscopy (EM)-labeling strategy and zero-shot deep-learning-based tracing with high-resolution volume EM to reconstruct growth cones of identified callosal projection neurons in the developing mouse brain. Mediolaterally extending growth cones exhibited a planar, fan-shaped morphology, preferentially expanded along the dorsoventral axis. Filopodia extended from the edges of this plane and aligned predominantly with the same axis but showed no directional bias toward the growing direction. Longitudinal in vivo imaging revealed that increased growth cone complexity associated with filopodial extension coincided with lower instantaneous velocity but preceded accelerated extension. Together, these findings reveal a dorsoventrally oriented planar growth cone architecture in which filopodial extension precedes accelerated axon growth, suggesting a structural constraint that may restrict axon navigation to the callosal plane.

## Main

The mammalian neocortex is characterized by precisely organized long-range axonal projections that preserve spatial relationships between cortical regions and their distant targets^4^. Among these, callosal projection neurons connect largely homotopic regions of the two cerebral hemispheres through the corpus callosum and thereby support bilateral integration of cortical information^1^. Abnormalities of the corpus callosum are observed in numerous human neurodevelopmental disorders^5,6^. During development, callosal axons cross the midline and, after entering the contralateral hemisphere, leave the white matter and extend radially through the cortex before terminal arborization. During these processes, axons navigate defined routes to reach specific targets within a densely packed environment composed of existing neurites, glial cells, and blood vessels^7^.

Extrinsic guidance cues recognized by the growth cone, a specialized, expanded structure at the tip of the extending axon, help direct precise axon targeting. Studies of axon guidance cues and their receptors on growth cones have suggested that growth cone structure shapes axonal growth behavior, prompting investigations of growth cone morphology in vivo in both invertebrates and vertebrates^2,3^. These studies have revealed that growth cone morphology varies across axonal tracts and locations, and live imaging has further shown that growth cone structure changes dynamically during axon extension and correlates with distinct modes of behavior^8–13^. In addition, previous electron microscopy (EM) studies have investigated growth cone ultrastructure at nanometer-scale resolution to understand the structural basis of cell-intrinsic growth cone regulation. Although these studies described filopodia and lamellipodia in growth cones, they have largely been limited to observations of two-dimensional (2D) thin sections or to non-mammalian species^8,14–17^. Moreover, dimensionality itself can influence growth cone morphology, as a recent study showed that growth cones in a 3D collagen matrix exhibit far fewer filopodia and greater microtubule extension into the leading region than those in 2D culture^18^. Thus, despite the importance of growth cone structure in directing axon navigation, how growth cones are organized ultrastructurally in 3D during navigation within the developing mammalian brain remains poorly understood.

Volume electron microscopy (vEM) is a promising approach for investigating the three-dimensional ultrastructure of growth cones in the brain, as advances in automated serial scanning electron microscopy (sSEM) have enabled imaging of increasingly large tissue volumes at nanometer-scale resolution. However, because these volumes remain small relative to the size of the brain, the neuronal subtypes from which analyzed axons originate cannot be identified in most cases. In addition, identifying rare and small structures such as growth cones in densely packed brain tissue is a major challenge in vEM datasets. While photoconversion of fluorescent-dye injected into regions containing neurons of interest has been used to identify growth cones by EM^19,20^, this approach requires intense illumination and can therefore be limited by light penetration in thick tissue. In contrast, genetically encoded peroxidases, including HRP and APEX2, catalyze diaminobenzidine (DAB) deposition independently of illumination^21,22^, facilitating labeling of genetically defined structures throughout large tissue volumes. This approach can therefore link axonal ultrastructure in vEM datasets to defined neuronal populations and facilitate the identification of rare structures such as growth cones. However, peroxidase activity can potentially perturb cellular physiology or obscure intracellular ultrastructure^22,23^.

A further challenge is tracing axons to their growth cones and reconstructing their 3D structures within densely packed membrane structures, tasks that have largely been performed manually^24,25^. Deep-learning-based analyses have enabled reconstruction of cortical networks and characterization of neuronal ultrastructure from large vEM datasets^26–28^. However, these approaches still require substantial manual effort to generate training datasets and proofread automated segmentations. Although zero-shot segmentation models such as the Segment Anything Model (SAM^29^) can segment structures on which they were not specifically trained and thereby eliminate the need for task-specific training data, they still require the user to supply a prompt in each section, typically a box drawn around the target, to specify which axon to follow. Consequently, tracing axons across hundreds of serial sections with these models still requires substantial manual labor.

In this study, we determine the 3D nanoscale ultrastructure of callosal projection neuron growth cones in the mouse brain using vEM datasets acquired by serial block-face scanning electron microscopy (SBF-SEM). We developed a genetically encoded tag that labels the extracellular space surrounding specific neurons with DAB, enabling growth cones to be identified within large-volume SBF-SEM datasets. We further developed an automated tracing approach using the promptable segmentation foundation model SAM, which segments the structure indicated by a user-supplied box, without task-specific training, allowing labeled axons to be traced and the 3D structures of dozens of growth cones to be reconstructed in the brain. We revealed that growth cones exhibited a planar, fan-like morphology preferentially oriented along the dorsoventral axis both while crossing the brain midline and after crossing it, but had almost no filopodia during midline crossing and multiple filopodia after crossing. In vivo two-photon imaging of the early postnatal mouse brain further revealed that a transient decrease in growth cone circularity preceded accelerated extension. By combining nanoscale 3D reconstruction with quantitative analysis of growth cone dynamics in the mammalian brain, our study reveals that growth cone morphology of callosal projection neurons is spatially polarized along the dorsoventral axis and undergoes dynamic changes that precede accelerated axon extension.

### Extracellularly expressed HaloTag enables identification of defined neurons in vEM datasets

To identify growth cones of callosal projection neurons in volume EM datasets, we first established a method to label specific neurons in grayscale vEM images by introducing contrast into the extracellular space, which provides a homogeneous and non-obstructive labeling target^30^. Although genetically encoded peroxidase-mediated DAB labeling has been widely used to label specific structures both in vitro and in vivo^30–36^, previous studies have shown that ectopic peroxidase expression can induce aberrant intracellular structures^22,23^. We therefore expressed a GPI-anchored HaloTag on the outer leaflet of the plasma membrane in living neurons and applied a horseradish peroxidase (HRP)-conjugated HaloTag ligand after tissue fixation **(Fig. 1a)**.

**Figure 1.**
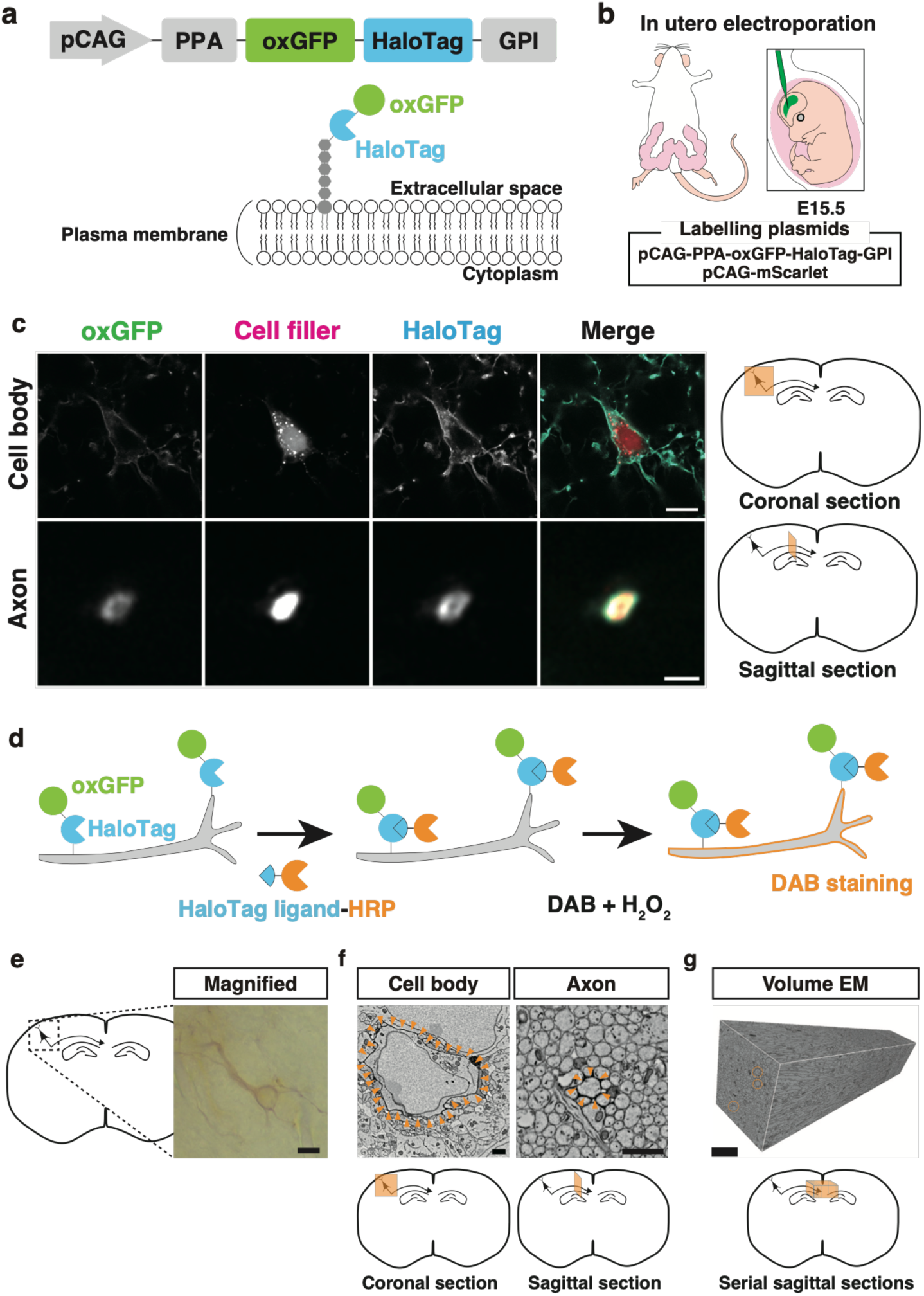
Extracellularly displayed HaloTag enables genetic labelling of specific neurons for electron microscopy. **a**, Schematic of the pCAG-PPA-oxGFP-HaloTag-GPI construct and membrane localization of the encoded oxGFP-HaloTag fusion protein through a GPI anchor. **b**, Schematic of in utero electroporation at E15.5 with pCAG-PPA-oxGFP-HaloTag-GPI and pCAG-mScarlet plasmids. **c**, Representative fluorescence images of the cell body and axon of labeled callosal projection neurons. Scale bars, 10 μm (cell body) and 1 μm (axon). **d**, Schematic of HaloTag-based HRP labeling followed by DAB staining. **e**, Representative bright-field image of a DAB-stained neurons in a coronal brain section. Scale bar, 10 μm. **f**, Representative electron micrographs of a labeled neuronal cell body and axon cross-section. Orange arrowheads indicate electron-dense DAB reaction products along the plasma membrane. Scale bars, 1 µm. **g**, Representative 3D reconstruction of a volume EM dataset acquired from a sagittal section. Orange circles indicate labeled axons. Scale bar, 5 µm.

To implement this membrane-targeted labeling strategy for both light and electron microscopy, we expressed a fusion protein, hereafter termed exHalo, consisting of oxGFP, a GFP variant optimized for oxidizing environments^37^, HaloTag, and a GPI-mPrion anchor in callosal projection neurons in the perinatal mouse neocortex **(Fig. 1a,b)**. mScarlet was co-expressed as a cytoplasmic cell filler to visualize the entire cell and facilitate comparison with the membrane localization of exHalo. To verify membrane localization and HaloTag ligand accessibility, brains were fixed at postnatal day 2 (P2) and sectioned into 100-µm-thick slices. Confocal fluorescence imaging showed that the native oxGFP signal sharply outlined the plasma membrane of both somata and axons, consistent with the expected distribution of the GPI-anchored protein, whereas the co-expressed mScarlet cell filler labeled the cytoplasm **(Fig. 1c)**. Treatment with the HaloTag SaraFluor 650T ligand produced a labeling pattern similar to that of oxGFP, confirming localization of HaloTag on the extracellular surface of the plasma membrane and demonstrating that it retained ligand-binding activity after aldehyde fixation. To label exHalo-expressing neurons for EM, brain slices were treated with an HRP-conjugated HaloTag ligand (Halo-HRP). Following incubation with Halo-HRP, we performed a biotinylated tyramine (BT)-glucose oxidase (GO) reaction, which induced local deposition of biotin around Halo-HRP-labeled sites^38^. The deposited biotin then recruited the avidin-biotin-peroxidase complex (ABC), thereby increasing the local peroxidase activity available to catalyze subsequent DAB polymerization **(Fig. 1d)**. This amplification procedure produced clear labeling with minimal background, as visualized by bright-field microscopy **(Fig. 1e)**. SEM imaging of ultrathin sections from DAB-stained brain slices confirmed that electron-dense DAB precipitates were confined to the extracellular surface of the plasma membrane, whereas the intracellular space remained free of dense labeling, preserving the visibility of internal organelles and ultrastructural details in both somata and axons **(Fig. 1f)**.

To investigate the 3D ultrastructure of callosal axons during development, we used SBF-SEM to acquire serial EM images of developing mouse neocortices *in utero* electroporated unilaterally at E15.5 in order to label layer 2/3 cortical pyramidal neurons (L2/3 CPNs) with exHalo and mScarlet constructs. To maximize the likelihood of capturing exHalo-labeled growth cones within the limited imaging volume, we selected sagittal sections from regions expected to contain abundant growth cones. Sections were first examined by confocal microscopy, and those showing the largest change in fluorescence signal between adjacent sections were selected for subsequent SBF-SEM imaging, as these were expected to contain a high density of axon terminals. By optimizing resin preparation, we obtained serial 2,400,000 µm square large sections at a resolution of 5–7 nm/pixel, with slicing steps of 25–40 nm **(Fig. 1g)**. In each slice, approximately ten regions of extracellular DAB staining corresponding to individual labeled axons were readily identified.

### Automated prompt propagation enables zero-shot tracing of axons in volume EM datasets

Identifying growth cones of labeled neurons requires tracing axonal shafts across hundreds of sections to their distal terminals. Although promptable zero-shot deep-learning models such as SAM greatly reduce the manual labor required for image segmentation, manually drawing bounding-box prompts on each section to specify the axon to be followed remains a major rate-limiting step. We therefore developed a SAM-based segmentation framework termed CLAMP (Closed-Loop Auto-Mask Propagation), in which the user draws a single bounding box on a seed section and subsequent prompts are generated automatically from the preceding segmentation mask. Within each section, CLAMP iteratively recomputes the bounding box and re-runs SAM until the segmentation converges, allowing misaligned prompts that degrade the initial one-pass SAM mask to be corrected within a few iterations (**Fig. 2a,b**). Closed-loop refinement also increased tolerance to prompt misplacement, maintaining greater overlap with the segmentation obtained from an undisplaced prompt than did a single SAM inference (**Extended Data Fig. 1**).

**Figure 2.**
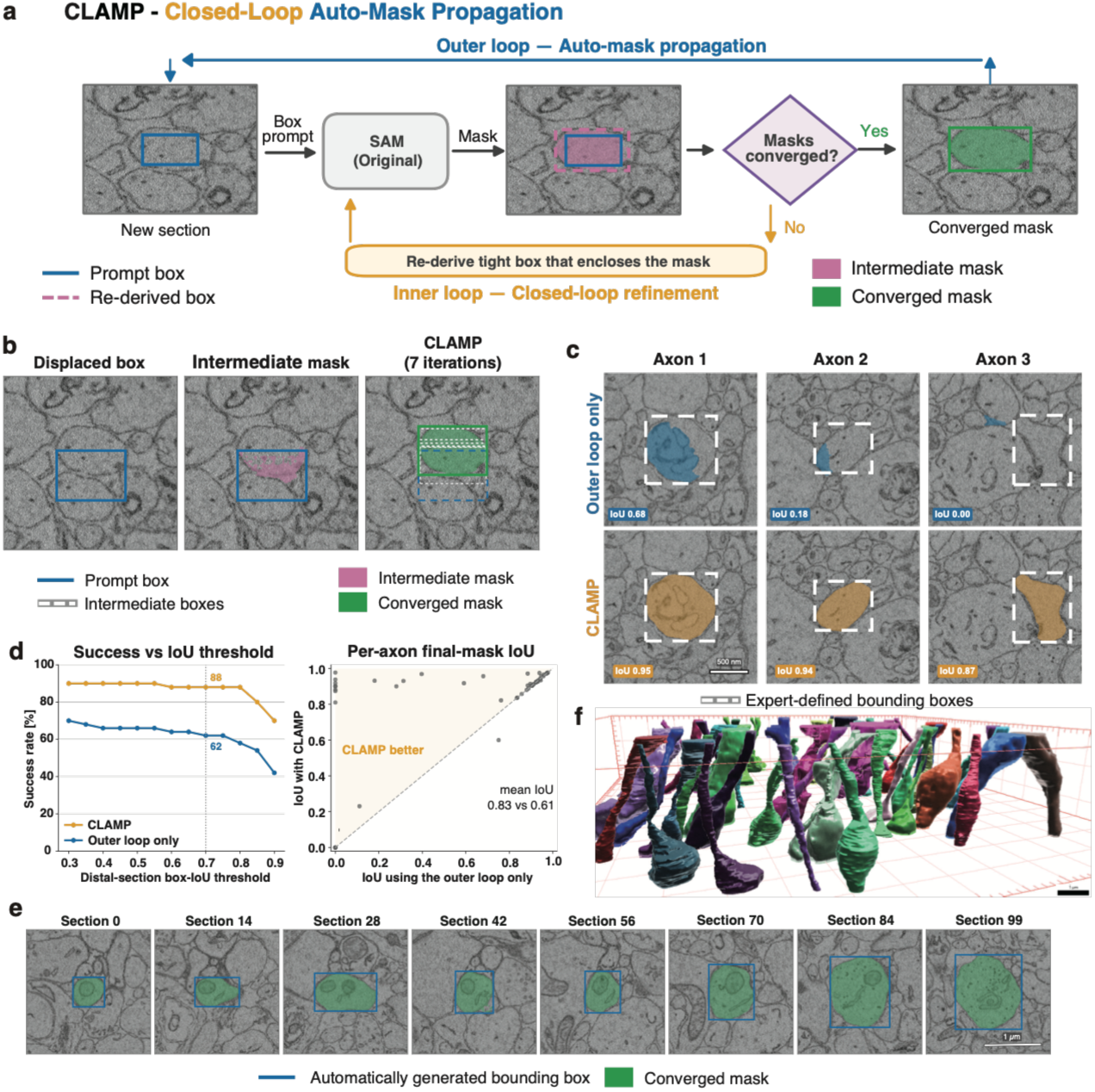
Closed-loop prompt refinement enables zero-shot tracing of axons through serial-section EM. **a**, The CLAMP tracing loop. A SAM mask is converted into a tight bounding box (Re-derived box) that prompts SAM again within the same section iteratively until masks converge, and the converged mask supplies the prompt box for the next section. The prompt box (blue), intermediate mask (magenta), re-derived box (magenta, dashed) and converged mask (green) are indicated. SAM denotes the original Segment Anything Model (ViT-H), used without training or fine-tuning. **b,** Example of prompt re-anchoring after deliberate displacement of the prompt box by 35% of the box side. The displaced prompt box (left) produces only a partial mask on the first pass (middle). During seven iterations of closed-loop refinement, the re-derived boxes progressively re-anchor to the axon profile, yielding the converged mask (right). Blue, prompt box; white dashed lines, intermediate boxes; magenta, intermediate mask; green, converged mask and its bounding box. **c,** Distal-section outcomes for three axons. With the outer loop alone, the mask cannot re-expand once it has shrunk, resulting in incomplete coverage of the distal profile for Axons 1 and 2 and the mask slipping off the target for Axon 3 (top, blue). With CLAMP, the mask follows the axon profile in each case (bottom, orange). Dashed boxes indicate expert-defined bounding boxes on the final section; values indicate box IoU. All panels are at the same magnification. Scale bar, 500 nm. **d,** Left, percentage of the 50 benchmark axons meeting each distal-section box-IoU threshold with the outer loop alone (blue) or CLAMP (orange); dotted line indicates the primary threshold of 0.7. The CLAMP condition includes the membrane gate (Methods). Right, paired per-axon final-mask IoU obtained with CLAMP versus the outer loop alone. **e,** Serial-section montage of one traced axon showing the automatically generated bounding box (blue) and converged mask (green) at the indicated sections, all at the same magnification. Scale bar, 1 µm. **f,** Three-dimensional reconstructions of the 50 benchmark axons assembled from the per-section masks, with each axon shown in a different color. The depth axis is scaled threefold. Scale bar, 1 µm.

We benchmarked CLAMP on an SBF-SEM volume of P5 mouse corpus callosum by providing a single bounding-box prompt for each of 50 unlabeled axons and allowing automatic tracing across 100 serial sections. Segmentation accuracy was assessed at the distal endpoint against an expert-drawn ground-truth box, with correct tracing defined as a distal-section box intersection over union (IoU) ≥ 0.7 (Methods). By this criterion, propagation without prompt correction correctly traced 31 of 50 axons, whereas CLAMP traced 44 of 50 (McNemar exact test, *P* = 1 × 10^-3^), with mean IoU increasing from 0.61 to 0.83 (Wilcoxon signed-rank test, *P*= 8 × 10^-5^) (**Fig. 2c,d**). Representative SBF-SEM sections show automatically generated boxes and SAM masks faithfully following each target axon through dense neuropil and adapting as the axonal profile enlarges from a thin cross-section to a much larger profile along the traced range (**Fig. 2e**). Three-dimensional reconstructions assembled from the per-section masks reproduced individual axon trajectories across the imaged volume (**Fig. 2f**).

### Developmental stage-dependent 3D ultrastructure of growth cones in the corpus callosum

Previous fluorescence microscopy studies have shown that the number of filopodia in a growth cone varies with its location and dynamic state^8,39–41^. We therefore first investigated by fluorescence microscopy whether neocortical growth cones show distinct morphologies depending on their location and developmental stage. Callosal projection neurons were labeled by expression of the green fluorescent protein mGreenLantern, and their growth cones were examined in coronal brain sections near the midline at P2 and within the corpus callosum and neocortex at P5 (**Extended Data Fig. 2a**). Consistent with previous studies reporting increased filopodial abundance on growth cones at axon guidance decision points^8^, growth cones located beyond the midline at P5 possessed significantly more filopodia-like protrusions than those located near the midline at P2 (**Extended Data Fig. 2b**). Because filopodia are thin structures, often less than 200 nm in diameter^42^, their number and morphology cannot be fully resolved by fluorescence microscopy. We therefore characterized growth cone morphology at these selected stages at higher spatial resolution using the vEM workflow established above.

To examine the ultrastructure of growth cones in P2 and P5 brains, DAB-labeled callosal axons identified in the SBF-SEM volumes were followed and segmented through serial sections to their distal endings using CLAMP. CLAMP-generated segmentations were manually corrected where necessary, and the axons were reconstructed in 3D. SBF-SEM datasets were acquired from multiple regions of individual specimens obtained from three P2 mice and one P5 mouse (**Extended Data Fig. 3a**). In total, 27 (for P2) and 11 (for P5) growth cones were reconstructed for subsequent morphological analysis (**Extended Data Fig. 3b,c**). Consistent with the fluorescence images 3D-reconstructed axon terminals corresponding to growth cones at P5 were significantly enlarged relative to control shaft regions and exhibited complex morphologies characterized by numerous protrusions **(Fig. 3a-f, Supplementary Video 1-4)**. More than 80% of P5 growth cones possessed at least one filopodium **(Fig. 3e)**. Callosal axon terminals at P2 were also enlarged relative to the axon shaft **(Fig. 3f)**, however, even at nanometer-scale EM resolution, most P2 growth cones lacked detectable filopodial protrusions **(Fig. 3e).** These results indicate a stage-specific morphological transition in callosal growth cones, in which filopodia are largely absent at P2 and become prominent by P5.

**Figure 3.**
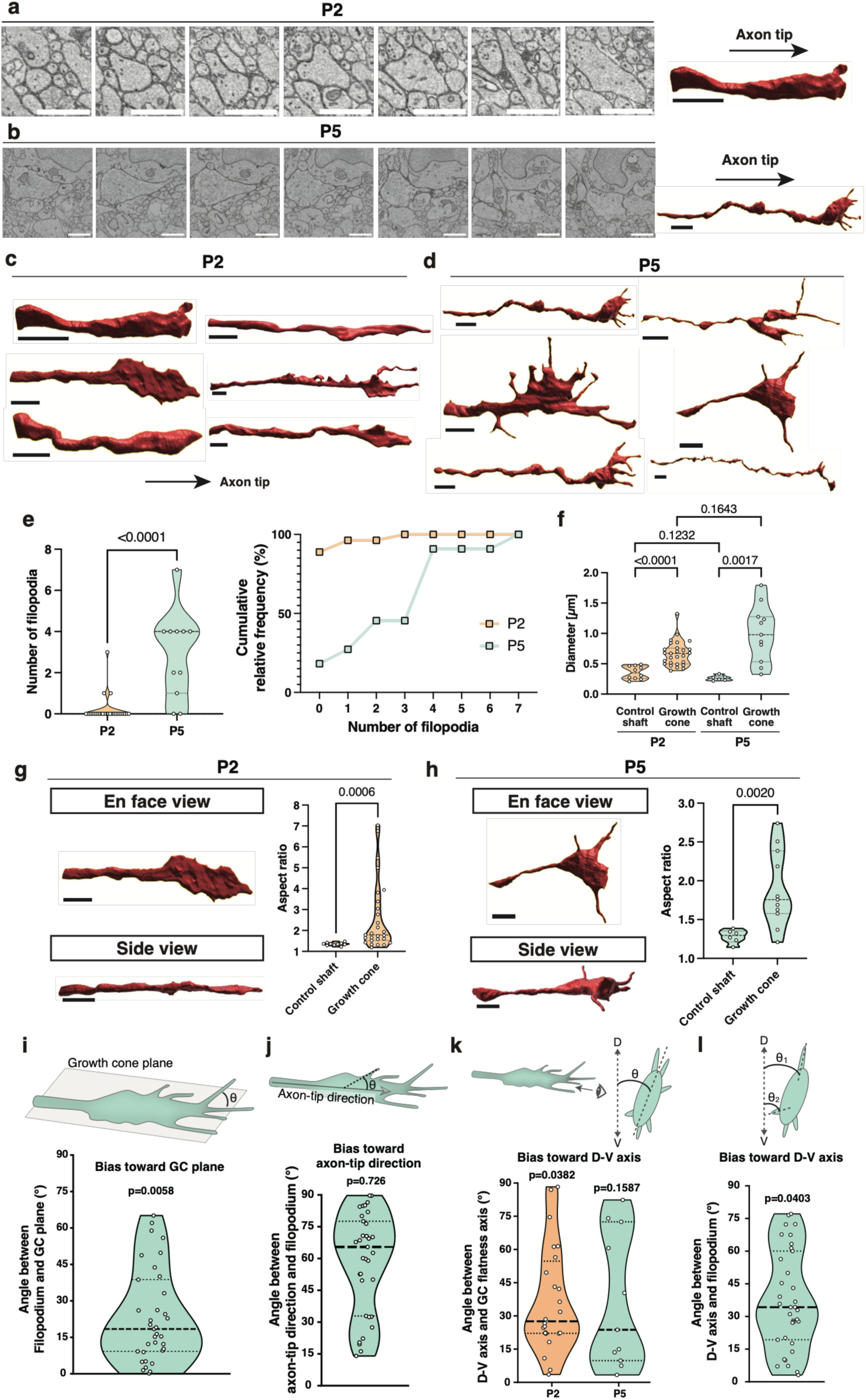
Neocortical growth cones are planar along the dorsoventral axis and extend filopodia from their edges. **a**–**d**, Representative serial EM images and 3D reconstructions of P2 (**a**) and P5 (**b**) growth cones, with additional reconstructions at P2 (**c**) and P5 (**d**). Arrows show axon-tip direction. **e**, Number of filopodia per growth cone (left) and cumulative frequency distributions (right). n = 27 growth cones from 3 mice at P2; n = 11 growth cones from 1 mouse at P5. **f**, Cross-section diameters of DAB-labled axon shafts (control shaft) and growth cones at P2 and P5 were compared. **g,h**, Representative en face and side views of reconstructed growth cones and width-to-thickness aspect ratios of control shafts and growth cones at P2 (**g**) and P5 (**h**). **i,j**, Acute angles of P5 filopodia relative to the growth cone plane (**i**) and to the axon-tip direction within the plane (**j**). For i, 0° was defined as extension within the growth cone plane and for j, 0° was defined as alignment with the axon-tip direction. **k,l**, Acute angles between the neocortical dorsoventral (D–V) axis and the long axis of P2 and P5 growth cone plane (**k**) or P5 filopodia (**l**). Scale bars, 2 µm (**a–d**,**g**,**h**).

### Neocortical growth cones and edge-derived filopodia form a dorsoventrally aligned plane

Although ultrastructural analyses of growth cones have been performed in vitro, growth cone morphology has been shown to depend highly on the culture environment^18^. To investigate growth cone geometry in vivo, we first quantified the local cross-sectional anisotropy of 3D-reconstructed DAB-labeled axons in planes orthogonal to the axonal centerline. For each axon or growth cone, the mean major-to-minor axis ratio (aspect ratio) across serial cross-sections was treated as a single independent measurement. Whereas axonal shafts had mean per-axon aspect ratios of 1.35 at P2 and 1.28 at P5, indicating nearly circular cross-sections, growth cones showed substantially higher aspect ratios of 2.62 at P2 and 1.90 at P5, respectively **(Fig. 3g,h)**. These results indicate that growth cones adopt a planar rather than cylindrical geometry in the developing corpus callosum.

Because growth cone filopodia have been proposed to serve sensory roles in detecting guidance cues and other extracellular signals^43^, their spatial orientation may influence how growth cones sense their local environment. Visual inspection of 3D-reconstructed growth cones at P5 revealed that filopodia rarely originated from their broad, flat surfaces and instead emerged preferentially from their edges **(Fig. 3d)**. To quantify this spatial bias, the extension axis of each reconstructed filopodium was measured relative to the plane of the corresponding growth cone, with 0° representing extension within the growth cone plane and 90° representing extension orthogonal to this plane **(Fig. 3i)**. Strikingly, filopodial extension angles were significantly concentrated around 0° by a Rayleigh V-test, indicating that filopodia preferentially extended within the plane of the growth cone rather than uniformly in three dimensions **(Fig. 3i)**. Within this plane, however, filopodial directions showed no significant bias toward the axon-tip direction **(Fig. 3j)**.

Next, we examined whether the long axes of individual growth cone planes were randomly oriented within the tissue or preferentially aligned with the body axes. The orientation of each growth cone’s long axis was therefore expressed relative to the dorsoventral direction, defined as 0°. Rayleigh V-test analysis revealed that growth cone orientations were significantly biased toward the dorsoventral axis at P2, with a weaker but still apparent tendency toward the same orientation at P5 **(Fig. 3k)**. Consistent with this spatial organization, filopodial orientations were also significantly aligned with the dorsoventral axis at P5 **(Fig. 3l)**. These results indicate that growth cones adopt a planar, rather than radially symmetric, morphology that is preferentially aligned with the dorsoventral axis in the developing brain. Given the sensory roles of growth cone filopodia, this spatial restriction of filopodial extension may constrain growth cone sensitivity to cues within the local dorsoventral plane, while reducing sensitivity to cues in adjacent anteroposterior planes.

### Ultrastructural environment surrounding the growth cone

vEM provides comprehensive information about the membrane architecture of diverse cell types within an imaged volume. Taking advantage of this feature, we next examined the local cellular environment surrounding growth cones in vivo. We reconstructed cellular structures around a representative callosal growth cone and found that the growth cone and its filopodia were embedded within a complex arrangement of neighboring cellular profiles **(Extended Data Fig. 4)**. Notably, a neighboring cellular structure, likely an axonal shaft, was positioned at the bases of two flanking filopodia **(Extended Data Fig. 4, blue)**. This observation suggests a possible association between filopodial extension and local cell–cell interactions. Although some neighboring profiles were morphologically consistent with glial cells **(Extended Data Fig. 4, green)**, their cellular identities could not be determined unambiguously from ultrastructure alone. These observations suggest that local cellular architecture may influence the spatial trajectories of growth cone filopodia in vivo.

### In vivo two-photon imaging captures growth cone dynamics in the brain

Whereas previous studies have shown that growth cone morphology differs across neuronal subtypes^11^ and developmental stages^8,41^, our neuronal subtype-specific analysis at P5 revealed substantial morphological variability even among growth cones of the same neuronal subtype at the same developmental stage. This variability prompted us to ask whether growth cone morphology is associated with their behavior. To address this question, we established a time-lapse two-photon imaging approach that enabled longitudinal imaging of individual growth cones of callosal axons within the corpus callosum. Because dense axon fasciculation and frequent axonal crossings in the brain precluded reliable tracking of individual growth cones over prolonged periods, we sparsely labeled individual L2/3 callosal projection neurons with tdTomato using the Supernova system^44,45^ delivered by in utero electroporation at E15.5 and imaged on the contralateral white matter at P4 (**Fig. 4a**). Imaging growing axons in deep brain regions during early postnatal development was further limited by poor light penetration due to the high density of blood vessels and lipid content. We therefore empirically optimized the depth of cortical aspiration by repeatedly assessing growth cone dynamics at different depths (**Fig. 4b**). We also minimized the size of the cranial window (**Fig. 4c**), because larger openings caused the cortical surface to bulge outward owing to intracranial pressure. To permit prolonged imaging while repeatedly changing the z-position of the objective lens, we installed a custom 3D-printed imaging cup that maintained a stable reservoir of immersion water between the cranial window and the objective lens (**Fig. 4d**). To avoid analyzing axons that might have been damaged during cortical aspiration, axons located close to the aspirated surface were excluded from imaging (**Fig. 4b,e)**. Together, these technical optimizations enabled imaging of growing axons in the white matter beneath the primary somatosensory cortex at 90-s intervals for 1.5-3.5 h (**Fig. 4e,f**). These time-lapse recordings revealed that individual growth cones dynamically changed their morphology and growth speed over time (**Fig. 4f, Supplementary Video 5**).

**Figure 4.**
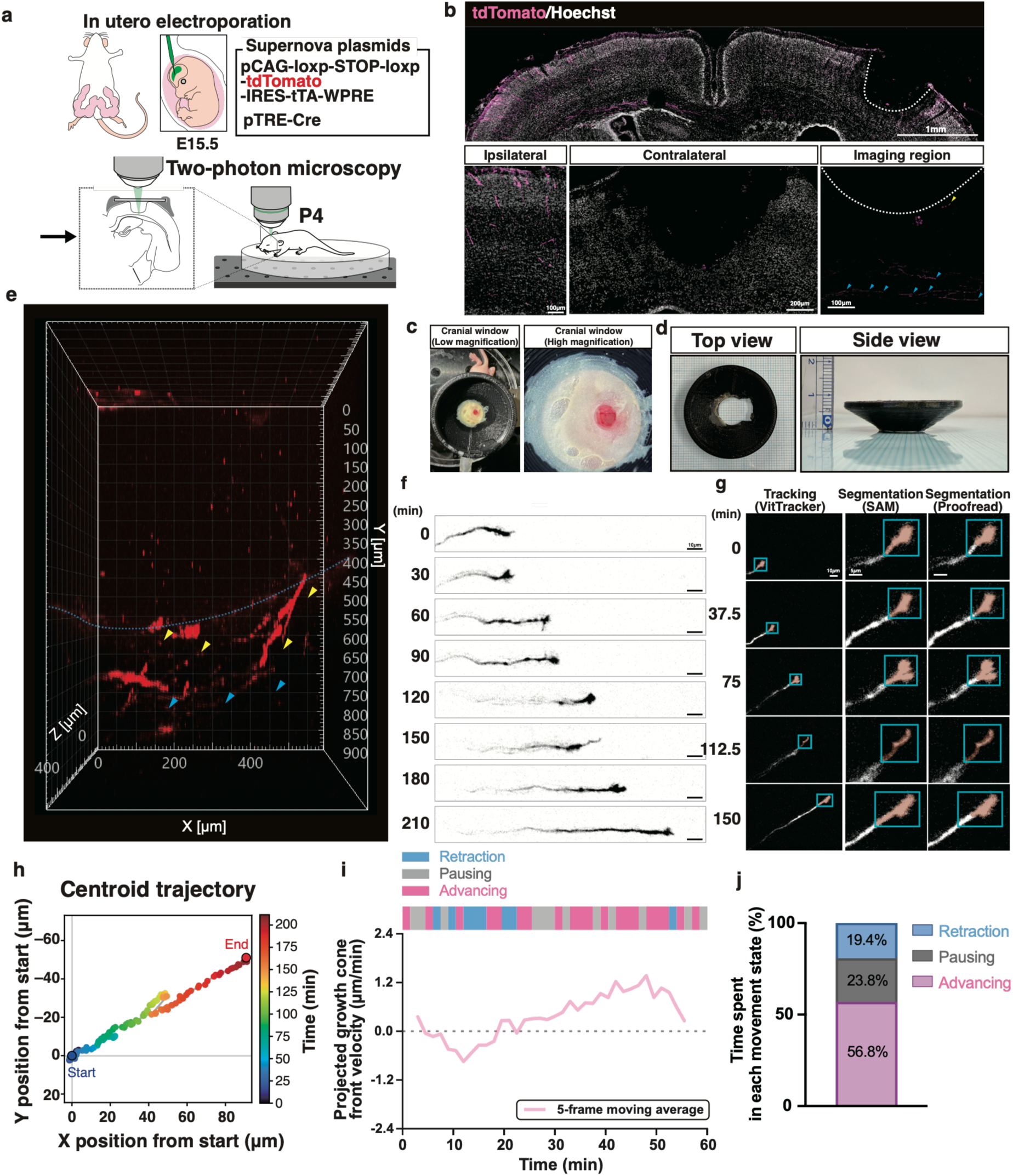
In vivo two-photon imaging of neonatal mice captures growth cone dynamics in the white matter. **a**, Schematic of the in vivo two-photon imaging setup for growing callosal axons in the neonatal mouse white matter. **b**, Representative coronal brain section showing growing callosal axons on the ipsilateral (electroporated) side and the cortical aspiration site (dotted line) on the contralateral side (top). Higher-magnification views of the ipsilateral cortex, and of the contralateral cortex and the aspirated region in separate sections from the same animal (bottom). Blue arrowheads, axons selected for imaging; yellow arrowhead, an axon excluded owing to its proximity to the aspirated surface. Scale bars, 1 mm (top), 100 μm (bottom left), 200 μm (bottom middle) and 100 μm (bottom right). **c**, Representative low– and high-magnification views of the cranial window. **d**, Top and side views of the custom-made 3D-printed imaging cup. **e**, Representative 3D view of an imaging volume extending from the brain surface to the white matter, showing growing axons within the white matter. Blue and yellow arrowheads indicate axons that were suitable and unsuitable, respectively, for quantitative analysis based on their distance from the aspirated surface. Dashed line: cortical surface after aspiration. **f**, Representative time-lapse images of a growing callosal axon in the white matter at P4. Scale bars, 10 μm. **g**, Representative images showing axon tracking using VitTracker, growth cone segmentation using SAM, and the final growth cone masks after manual proofreading. Time is shown relative to the first displayed frame. Scale bars, 10 μm (tracking) and 5 μm (segmentation). **h**, Representative trajectory of a growth cone centroid, color-coded according to elapsed time. **i**, Temporal changes in the five-frame moving-average velocity of a representative growth cone (magenta line). The colored bars indicate growth cone states classified using unsmoothed frame-to-frame velocity: advancing (magenta; *v* > 0.30 μm/min), pausing (grey; −0.30 μm/min ≤ *v* ≤ 0.30 μm/min), and retracting (blue; *v* < −0.30 μm/min). **j**, Proportions of observation time spent in the advancing, pausing, and retracting states across axons.

Although growth cone dynamics have been extensively characterized qualitatively, only a limited number of studies have quantitatively analyzed growth cone behavior^46^. To quantitatively examine how growth cone morphology relates to ongoing and subsequent behavior, reliable and efficient segmentation of growth cones across this large time-lapse dataset was required. To address this technical challenge, we first applied VitTracker, a Vision Transformer (ViT)-based discriminative tracker implemented in OpenCV, which reliably tracked individual growth cones across consecutive frames of long-term in vivo imaging (Fig. **4g,h**). Bounding boxes generated by VitTracker were then used as prompts for SAM, enabling automatic segmentation of the growth cone in each frame (**Fig. 4g**). After SAM-based segmentation, all segmentation results were manually proofread (**Fig. 4g**).

The trailing boundary of the growth cone would ordinarily be identified as the position at which the broadened distal region sharply narrowed into the axon shaft. However, this boundary was sometimes difficult to identify when the growth cone tapered gradually and merged smoothly with the shaft. To quantify growth cone translocation while minimizing the influence of local morphological fluctuations and avoiding the need to explicitly define the potentially ambiguous trailing boundary, we defined growth cone position as the 90th percentile of mask-pixel coordinates projected onto the trajectory axis, corresponding to the point beyond which the leading 10% of mask pixels were located (**Extended Data Fig. 5**; Methods). Following growth cone collapse, a new growth cone typically re-formed from a proximal broadened region and subsequently resumed extension. Because this process appeared as a large rearward displacement in our positional measurements rather than ordinary retraction, intervals spanning collapse and growth cone reformation were excluded from the movement analyses. Growth cones switched among advancing, pausing, and retracting states on a timescale of minutes, as classified by velocity (**Fig. 4i**; advancing, v > 0.30 μm/min; pausing, −0.30 μm/min ≤ v ≤ 0.30 μm/min; retracting, v < −0.30 μm/min). Individual advancing, pausing, and retracting bouts lasted a median of 3.0 min (IQR, 1.5–4.5 min; n = 170 bouts), 1.5 min (IQR, 1.5–3.0 min; n = 131 bouts), and 1.5 min (IQR, 1.5–1.9 min; n = 112 bouts), respectively. These states accounted for 56.8 ± 5.2%, 23.8 ± 2.8%, and 19.4 ± 2.8% (mean ± SEM) of the observation time, respectively (**Fig. 4j**). These values are consistent with those previously reported in organotypic brain slice cultures from golden hamsters, supporting the validity of our newly established in vivo two-photon imaging approach^10^.

### Growth cone complexity primes axons for accelerated extension

Next, we investigated the relationships between specific morphological features of segmented growth cones and their translocation dynamics (**Fig. 5a**). Growth cone morphology at each time point (*t*) was quantified using metrics including circularity (Cir), eccentricity (Ecc), and area. We then systematically examined the relationships between these morphological parameters and growth cone translocation dynamics (**Extended Data Figs. 6 and 7**). As also observed in the P5 vEM reconstructions, growth cones in the longitudinal imaging exhibited diverse morphologies with variable numbers of filopodia (**Supplementary Video 5**). Consistent with previous observations linking relatively simple growth cone morphologies to axonal advance^9^, circularity at *t_0_* [Cir(*t_0_*)] showed a positive correlation with instantaneous growth cone velocity, *v_0_* (r = 0.25; **Fig. 5b and Extended Data Fig. 6**). This relationship is also in line with the lower number of filopodia observed in growth cones near the midline, a morphology expected to be associated with higher circularity (**Fig. 3e and Extended Data Fig. 6**), where callosal axons extend faster than in subtarget regions of the corpus callosum^10^.

**Figure 5.**
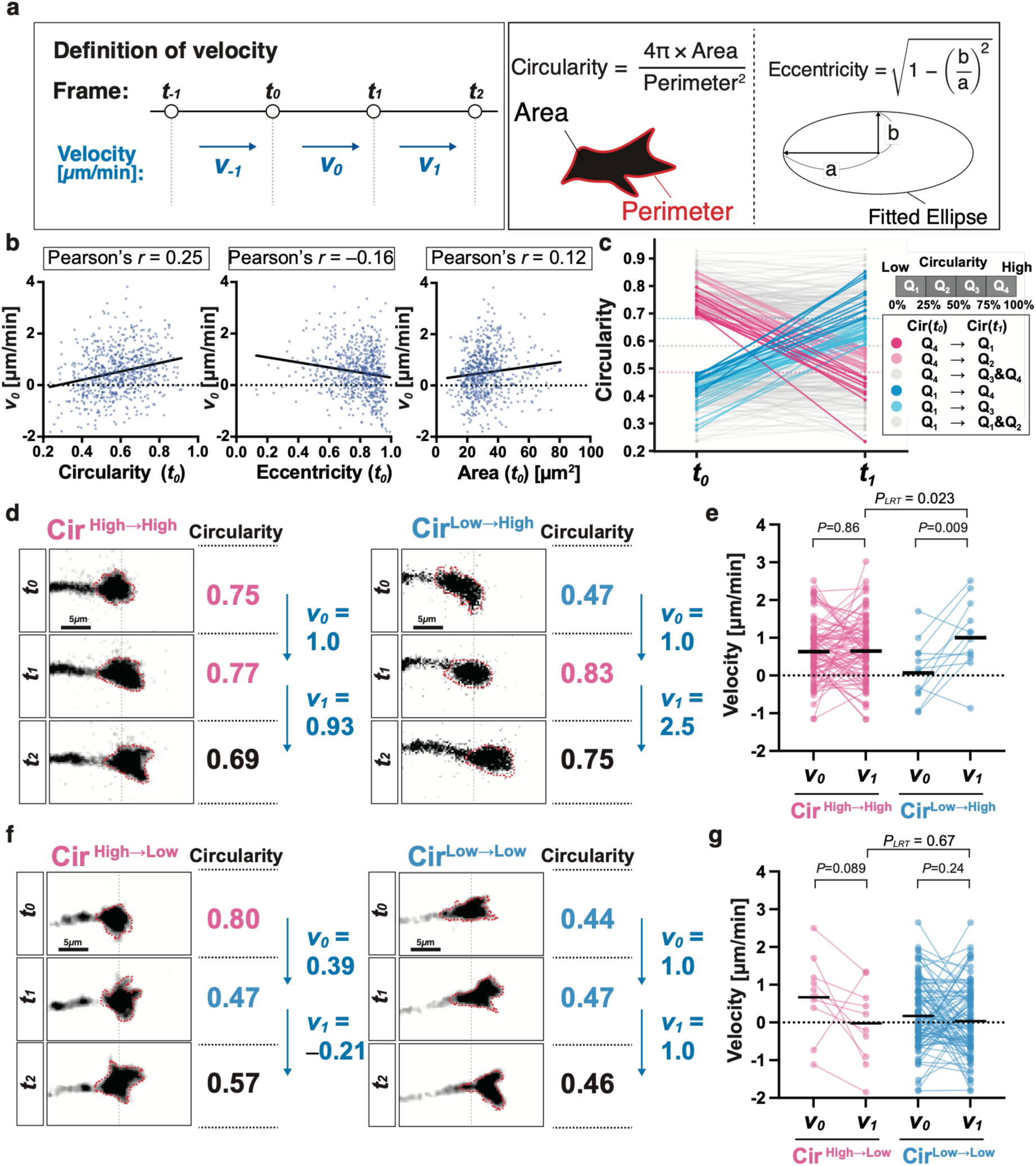
Growth cone complexity marks a state primed for accelerated extensiona, Velocity of growth cones were defined as follows: *v_i_,* velocity from *t_i_* to *t_i+1_* (μm/min), where Δ*t* represents the 1.5-min interval between consecutive imaging frames. Circularity, eccentricity and area were quantified from growth cone masks. Schematics illustrate the calculation of circularity and eccentricity. **b**, Association between growth cone morphology and velocity. Growth cone Cir(*t_0_*), Ecc(*t_0_*) and Area(*t_0_*) were plotted against *v_0_* (8 axons; *n* = 675 target time points). Target time points were included only when a complete sequence of four consecutive frames (*t_−1_* to *t_2_*) was available. Pearson’s correlation coefficients are shown. The corresponding 95% confidence intervals for circularity, eccentricity and area respectively, were 0.18 to 0.32, −0.23 to −0.081 and 0.044 to 0.19. **c**, Changes in circularity over one 1.5-min imaging interval among events starting in the lowest (Q_1_) or highest (Q_4_) quartile of Cir(*t_0_*). Quartile thresholds at (*t_0_*) and (*t_1_*) were calculated from all eligible events (n = 708). Dotted horizontal lines indicate the quartile boundaries. For events starting in Q_4_ of Cir(*t_0_*) (*n* = 177): Q_4_→Q_1_, 10 events (5.6%); Q_4_→Q_2_, 26 (14.7%); and Q_4_→Q_3_&Q_4_, 141 (79.7%). For events starting in Q_1_ (*n* = 177): Q_1_→Q_4_, 12 events (6.8%); Q_1_→Q_3_, 26 (14.7%); and Q_1_→Q_1_&Q_2_, 139 (78.5%). **d**, Representative time-lapse images of Cir^High→High^ and Cir^Low→High^ events. Images show the growth cone at *t_0_*, *t_1_* and *t_2_*. Scale bars, 5 µm. **e**, Changes in velocity for Cir^High→High^and Cir^Low→High^ events (Cir^High→High^, n=87 events from 8 axons; Cir^Low→High^, n=12 events from 6 axons). Horizontal bars indicate means. Within-group changes were assessed using paired t-tests. The between-group difference in *v_1_* was assessed using a linear mixed-effects model (LMM), with axon identity included as a random intercept. **f,** Representative time-lapse images of Cir^High→Low^ and Cir^Low→Low^ events, presented as in **d**. Scale bars, 5 µm. **g**, Changes in velocity for Cir^High→Low^ and Cir^Low→Low^ events (Cir^High→Low^, n = 10 events from 6 axons; Cir^Low→Low^, n = 94 events from 6 axons). Lines, horizontal bars and statistical analyses are as described in **e**. Dotted lines in d and f indicate growth cone boundaries defined as in Fig. 4.

Taking advantage of the high temporal resolution of the longitudinal imaging, we next investigated the relationship between morphological dynamics and subsequent velocity. Within a single 1.5-min imaging interval, more than 5% of growth cones in the highest (Q_4_) or lowest (Q_1_) quartile of Cir(*t_0_*) transitioned to the lowest or highest quartile of Cir(*t_1_*), respectively (**Fig. 5c)**. We next examined whether these dynamic morphological transitions were associated with *v_1_*. Consistent with the positive correlation between Cir(*t_0_*) and *v_0_*, growth cones in the Q_4_ group of Cir(*t_1_*) showed higher *v_1_* than those in the Q_1_ group, regardless of Cir(*t_0_*) (**Extended Data Fig. 8)**. Interestingly, within the Q_4_ of Cir(*t_1_*), growth cones that had been in the Q_1_ of Cir(*t_0_*) (Cir^Low→High^) showed significantly higher *v_1_* than those that had been in the Q_4_ (Cir^High→High^) (**Fig. 5d,e**; the linear mixed-effects model estimated β = 0.625 µm/min; 95% CI, 0.093–1.156; likelihood-ratio test, χ²(1) = 5.18, *P* = 0.023). Consistent with this result, growth cones in the Cir^Low→High^ group accelerated on average from 0.11 µm/min to 1.05 µm/min (paired t-test, *P* = 0.009), whereas no significant change was detected in the Cir^High→High^ group (0.68 to 0.69 µm/min, *P* = 0.86; **Fig. 5e**). The difference in *v_1_* between the Cir^Low→High^ and Cir^High→High^ groups remained statistically significant after accounting for differences in *v_0_* between the groups (β = 0.653 µm/min; 95% CI, 0.129–1.177; likelihood-ratio test, χ²(1) = 5.83, *P* = 0.016), suggesting that the greater acceleration in the Cir^Low→High^ group was not solely attributable to its lower *v_0_*. By contrast, among events remaining in Q_1_ or transitioned from Q_4_ to Q_1_ at *t_1_*, *v_1_* did not differ significantly between the Cir^Low→Low^ and Cir^High→Low^ groups (**Fig. 5f,g**; *P* = 0.67). These results suggest that low circularity established a state permissive for subsequent acceleration, whereas a transition to higher circularity allowed this state to be translated into increased velocity.

## Discussion

By combining genetically encoded extracellular labeling, volumetric EM and deep-learning-based image segmentation with longitudinal in vivo two-photon imaging, we resolved the 3D ultrastructure and dynamics of neocortical growth cones in the developing brain. Surprisingly, growth cones of callosal projection neurons were planar and preferentially aligned along the dorsoventral axis. They largely lacked filopodia near the midline but elaborated multiple filopodia after crossing into the contralateral hemisphere. Our longitudinal imaging further suggests that sustained growth cone complexity may establish a permissive state for acceleration, whereas the subsequent transition to a more circular morphology is associated with the ensuing increase in axon extension velocity.

Although mechanisms regulating the midline crossing of callosal axons have been extensively studied, how these axons are guided after crossing the midline, including how they preserve their anteroposterior coordinates during development, remains largely unknown^15,47–50^. Given that growth cone filopodia can localize guidance receptors and signaling machinery^51^, our findings suggest that the dorsoventral organization of growth cones and their filopodia may constrain where extracellular cues are sensed and thereby bias directional decisions both within the corpus callosum and as callosal axons transition into the neocortex. Importantly, in contrast to previous reports of forward-directed filopodia in vivo^52^, filopodial orientations within the growth cone plane showed no bias toward the axon-tip direction, suggesting that this organization is not simply a reflection of forward growth.

While our longitudinal imaging revealed a positive correlation between circularity and instantaneous velocity, consistent with the qualitative classifications reported in previous in vivo studies^9,13,53–55^, our data further showed that growth cone complexity preceded subsequent acceleration. Together with the lack of filopodial bias toward the axon-tip direction, this temporal sequence suggests that increased growth cone complexity may enhance sensitivity to local guidance cues before accelerated extension. Although the underlying mechanism remains unresolved, these findings indicate that growth cone morphology is organized in both space and time, providing a structural framework through which developing axons may interpret local guidance cues and regulate their advance.

Although genetically encoded peroxidases, including membrane-targeted HRP and APEX-based reporters, generate cell-specific contrast for EM^22,23,30^, high expression of organelle-targeted peroxidases can induce organelle aggregation or morphological perturbations^22,23^, and membrane-targeted HRP can undergo endocytosis and enter intracellular compartments^30^. Moreover, plasma-membrane-targeted APEX2 can generate DAB reaction product that extends into the cytoplasm and obscures ultrastructural features in fine neuronal processes^36^. We found that HaloTag retained its ligand-binding activity after glutaraldehyde fixation, allowing HRP to be conjugated to exHalo after tissue fixation and thereby avoiding peroxidase expression in living cells. Because exHalo efficiently presents HaloTag on the extracellular surface, DAB reaction product remained confined to the extracellular side of labeled plasma membranes. Thus, exHalo, a fusion protein developed in this study, expands the scope of vEM by enabling genetically defined neurons to be identified within vEM datasets while preserving their intracellular ultrastructure.

To scale volumetric and longitudinal analyses, we combined automatic prompt generation with the frozen foundation model SAM for growth cone tracking and segmentation. Whereas prompt propagation across serial images can accumulate drift^56–59^, CLAMP iteratively refines prompts based on segmentation outputs without additional training or auxiliary learned modules. Extending CLAMP to generate multiple prompts at branch points could facilitate segmentation of mature neuronal arbors. For time-lapse fluorescence imaging, we instead used VitTracker to generate prompts for visually distinctive growth cones. Both strategies were integrated into a single open-source napari plugin for SAM-based segmentation (see Methods and Code Availability).

The dorsoventrally polarized growth cone ultrastructure identified in this study provides a framework for understanding how growth cones orient within the three-dimensional body axes and for uncovering the mechanisms that shape axonal trajectories during bilateral circuit formation. This discovery opens new avenues for determining how growth cone ultrastructure contributes to axon guidance across neuronal populations and developmental stages.

**Extended Data Figure 1.**
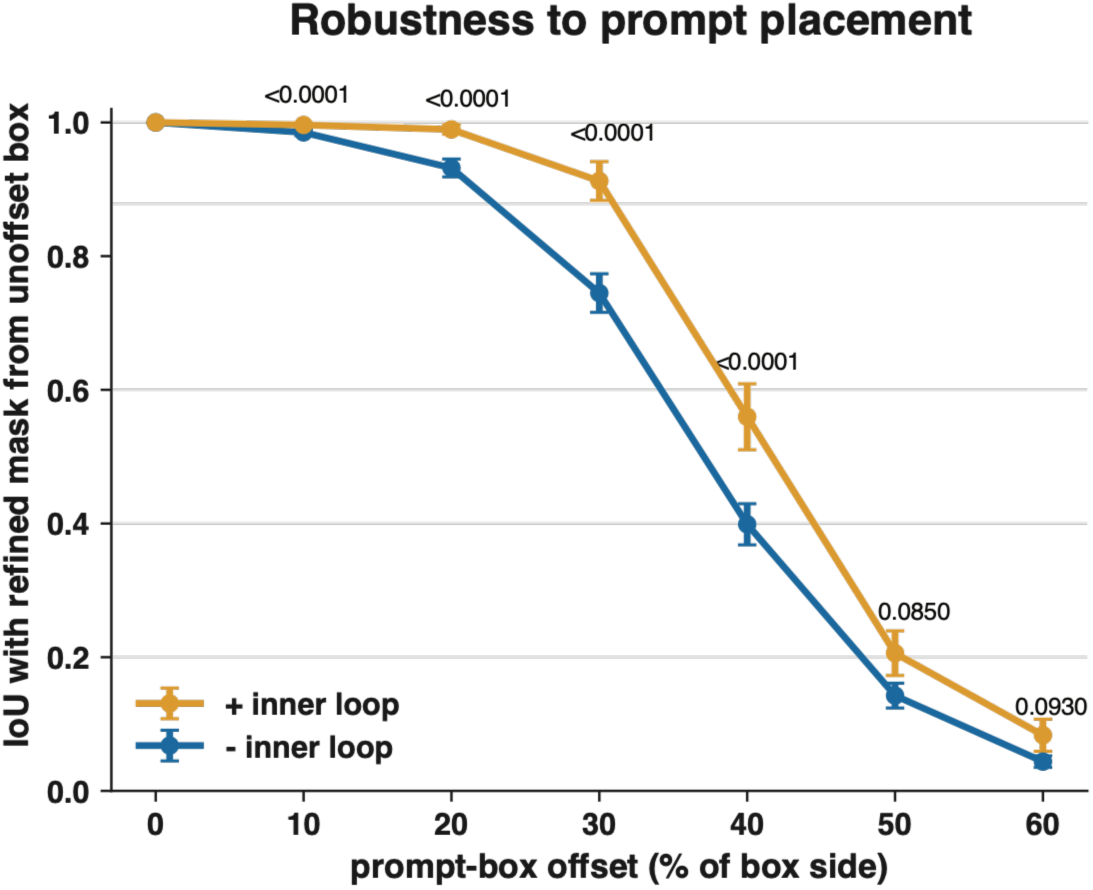
Closed-loop refinement increases robustness to prompt displacement. Axonal sections were re-segmented after shifting the prompt boxes in four directions by the indicated fraction of the box side length. Segmentation overlap was quantified as IoU with the mask obtained from the undisplaced prompt, comparing SAM run once (− inner loop, blue) with the refinement loop run to convergence in the same section (+ inner loop, orange). Mean ± s.e.m.; *n* = 24 sections. Two-sided Wilcoxon signed-rank test with Holm correction across the six displacement magnitudes (10 to 60 % in 10 % steps).

**Extended Data Figure 2.**
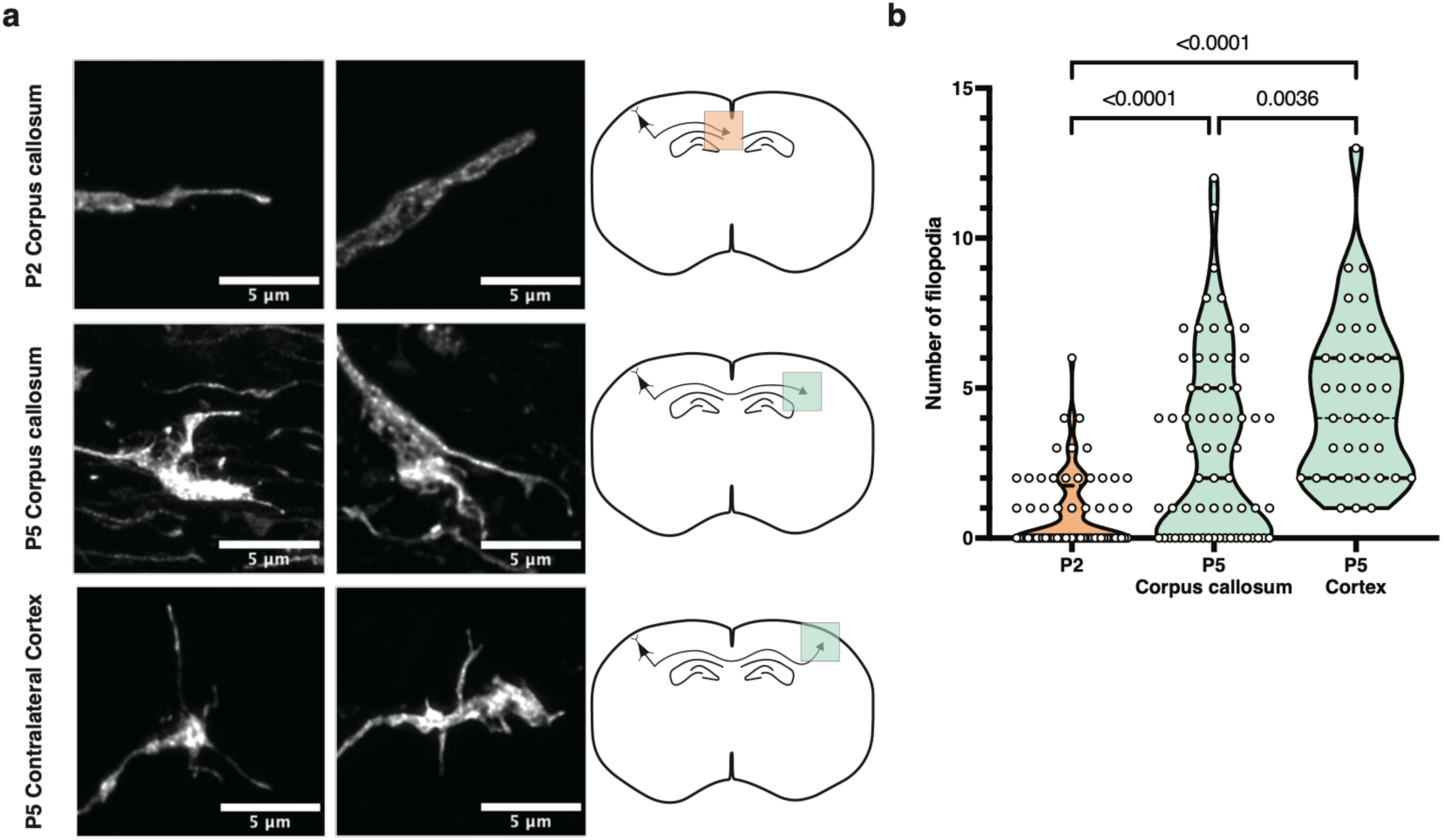
Confocal analysis of developmental and regional differences in growth cone filopodia. **a**, Callosal projection neurons were labeled by expressing the green fluorescent protein mGreenLantern. Representative confocal fluorescence images of their growth cones at P2 in the corpus callosum, at P5 in the corpus callosum and at P5 in the contralateral cortex are shown. Schematics indicate the imaging locations. Scale bars, 5 µm. **b,** Number of filopodia per growth cone at the indicated developmental stages and locations. *N* = 64, 65 and 37 growth cones, respectively. Kruskal–Wallis test.

**Extended Data Figure 3.**
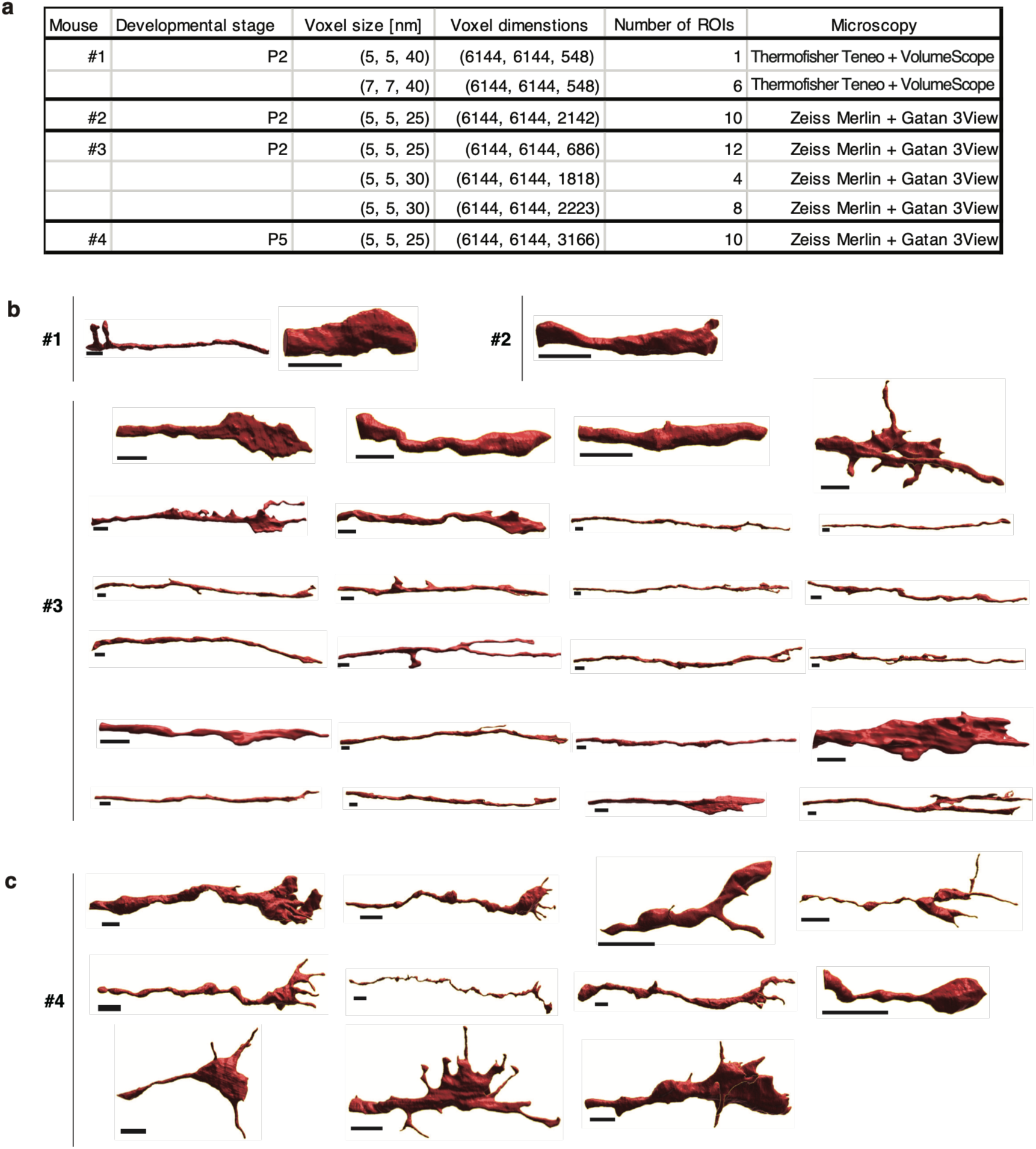
Overview of SBF-SEM datasets and 3D reconstructions of callosal growth cones. **a**, Summary of the SBF-SEM datasets acquired at P2 and P5, including voxel size, voxel dimensions and the number of independent imaging volumes. **b,** 3D surface reconstructions of all growth cones analyzed at P2. *n* = 27 growth cones from 3 mice. **c,** 3D surface reconstructions of all growth cones analyzed at P5. *n* = 11 growth cones from 1 mouse. Scale bars, 2 µm

**Extended Data Figure 4.**
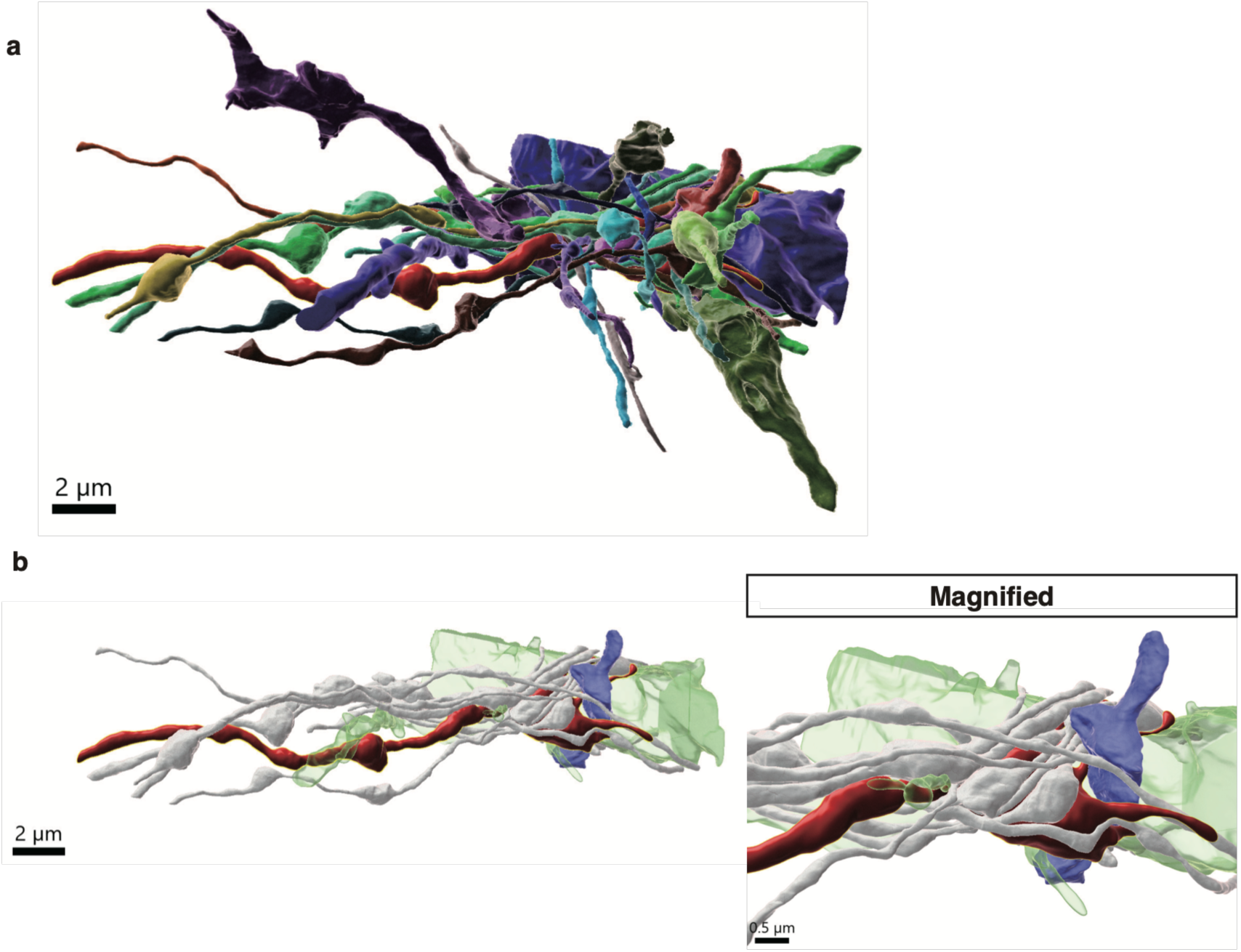
Dense reconstruction of the cellular environment surrounding a callosal growth cone. **a**, 3D surface reconstruction of a callosal growth cone and neighboring cellular structures within the imaged volume. Individual structures are shown in distinct colors. **b**, Rendering of a subset of the cellular structures shown in **a**: growth cone (red), most surrounding structures (white), and selected structures (green and blue). Right, magnified view of the region where two filopodia of the growth cone are closely adjacent to the cellular structure highlighted in blue. Scale bars, 2 µm (**a** and **b**, left) and 0.5 µm (**b**, right).

**Extended Data Figure 5.**
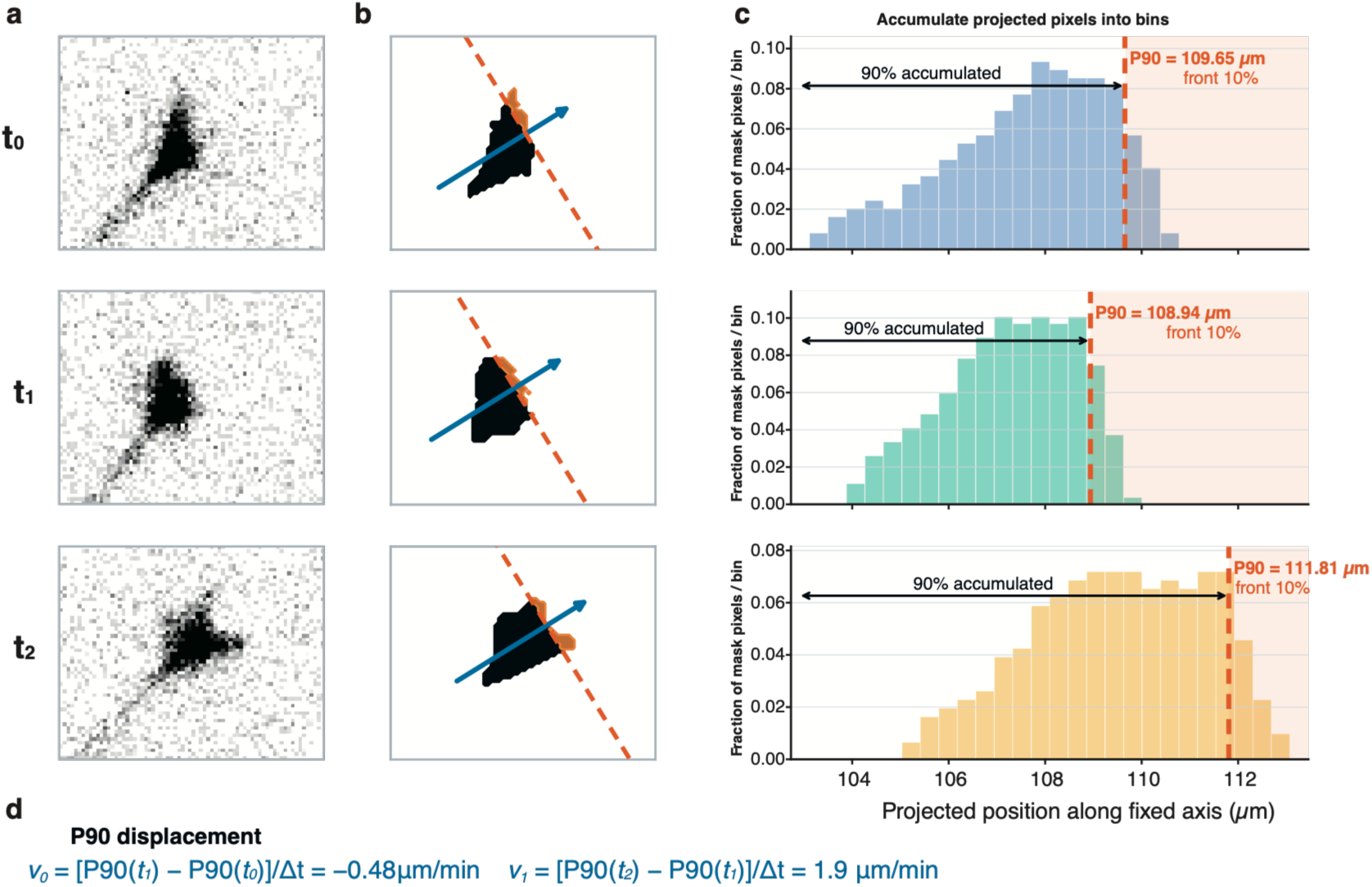
Calculation of the growth cone velocity and acceleration. **a**, Representative raw images of a growth cone at *t_0_*, *t_1_* and *t_2_*. **b,** Corresponding proofread binary masks and orthogonal projections of mask pixels onto a fixed global progression axis. The axis (blue arrow) was defined by the displacement from the first and the last growth cone centroids and held constant across all frames. The orange region contains the most anterior 10% of projected pixels, and the orange dashed line indicates the 90^th^-percentile position (P90). **c,** Distributions of projected mask-pixel positions, expressed as the fraction of total mask pixels in each bin. **d,** Calculation of P90-based velocities and acceleration.

**Extended Data Figure 6.**
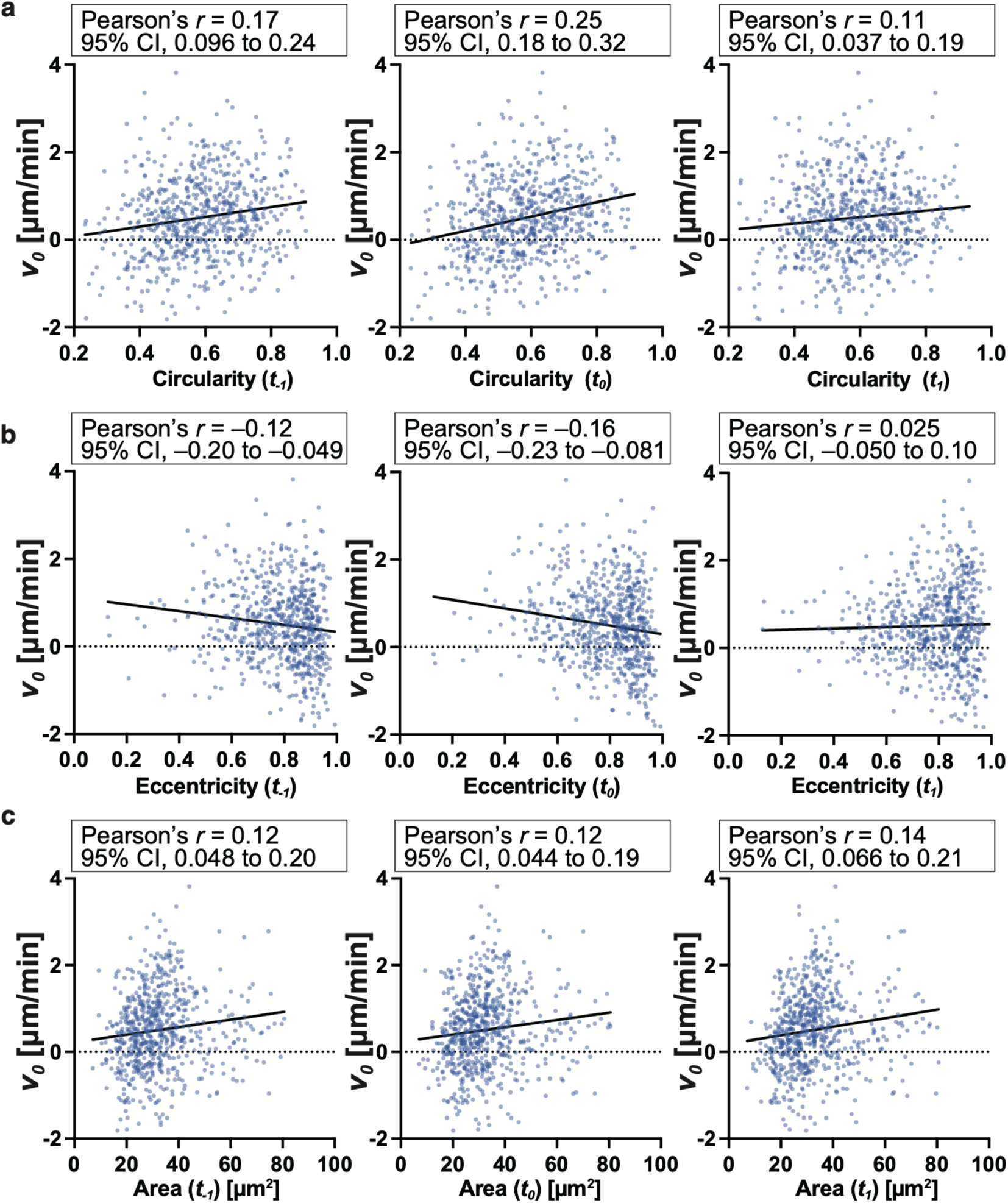
Associations between growth cone morphology at different time points and velocity. Circularity (**a**), eccentricity (**b**) and area (**c**) at *t*_−*1*_, *t_0_*, and *t_1_* were plotted against the *v_0_*. Each point represents one target time point, and solid lines indicate ordinary least-squares regression lines. Circularity and eccentricity are dimensionless, and area is expressed in μm^2^. Pearson’s *r* values and 95% confidence intervals are shown in the plots.

**Extended Data Figure 7.**
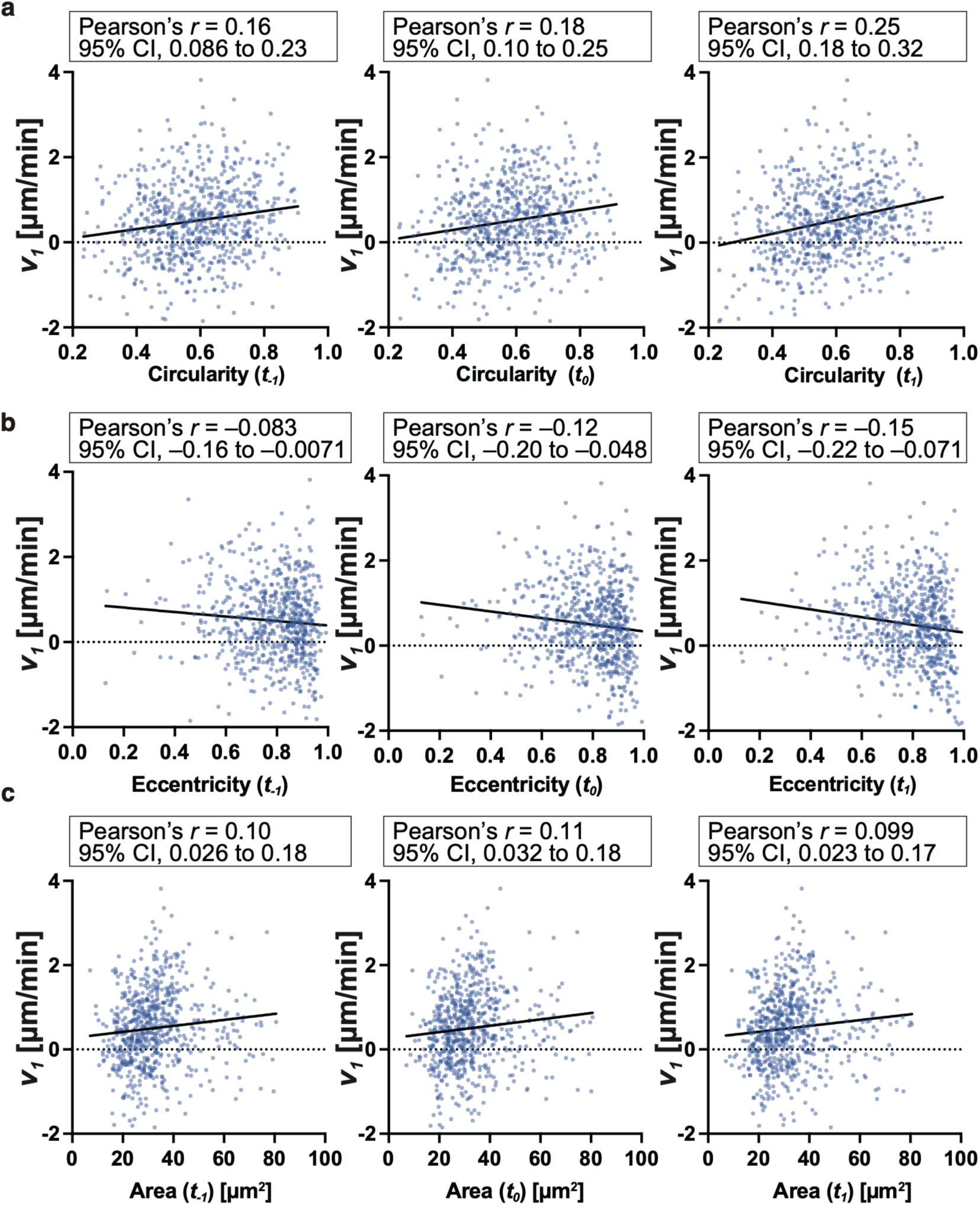
Associations between growth cone morphology at different time points and velocity. As in **Extended Data** Fig. 6, but with circularity (**a**), eccentricity (**b**) and area (**c**) plotted against *v*_1_.

**Extended Data Figure 8.**
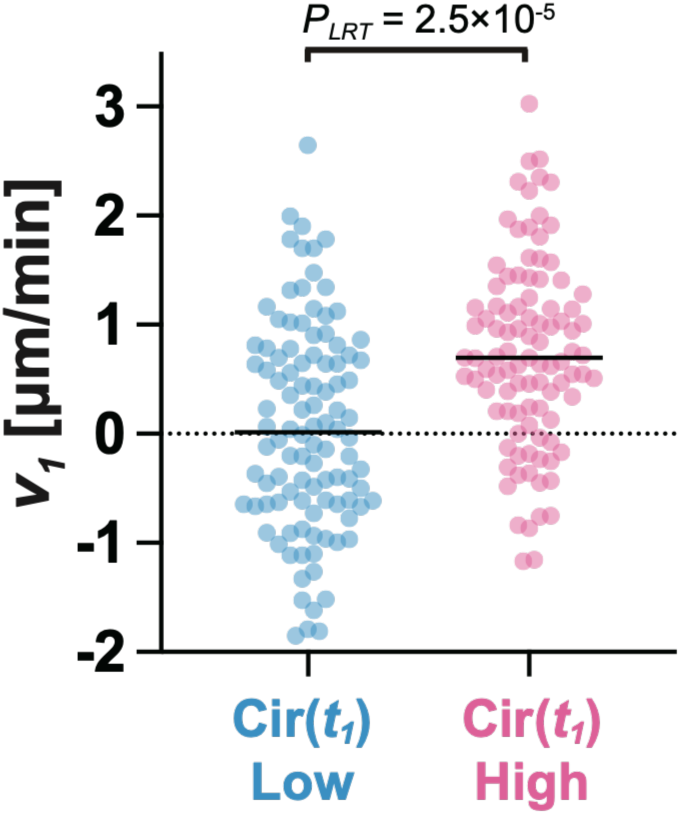
Growth cones that transitioned to high circularity showed higher concurrent velocity. Comparison of *v_1_* between Cir(*t_1_*)^Low^ and Cir(*t_1_*)^High^ events (Cir(*t_1_*)^Low^, n = 104; Cir(*t_1_*)^High^, n = 99). Horizontal bars indicate means. The linear mixed-effects models included axon identity as a random intercept. *P_LRT_* denotes the likelihood-ratio test of the Cir(*t_1_*) state term.

**Supplementary Video 1**. 3D reconstruction of a growth cone at P2.

SBF-SEM images with the segmentation and 3D reconstruction of a growth cone at P2.

**Supplementary Video 2.** 3D reconstruction of another growth cone at P2.

SBF-SEM images with the segmentation and 3D reconstruction of an additional growth cone at P2.

**Supplementary Video 3.** 3D reconstruction of a growth cone at P5.

SBF-SEM images with the segmentation and 3D reconstruction of a growth cone at P5.

**Supplementary Video 4.** 3D reconstruction of another growth cone at P5.

SBF-SEM images with the segmentation and 3D reconstruction of an additional growth cone at P5.

**Supplementary Video 5.** Longitudinal imaging of growing callosal axons in the P4 mouse brain

In vivo two-photon microscopy captures the dynamic morphological changes of advancing growth cones, with images acquired continuously at 90-s intervals for more than 3 h.

## Acknowledgements

The authors thank Drs. Megumu Takahashi and Yoko Ishida (Juntendo University) for plasmid preparation, Dr. Takumi Higaki for optimization serial image alignment, and Drs. Atsushi Miyawaki and Yusuke Niino (RIKEN CBS) for providing the plasmid containing GPI (mPrion) sequence, Atsuko Imai and Nobuko Hattori (National Institute of Physiological Sciences, Japan) for their technical assistance, Dr. Masayasu Taki for providing HaloTag ligand, and Drs Franck Polleux and Yukiko Gotoh for their critical reading of the manuscript.

This study was supported by JSPS KAKENHI (JP20K15894, JP22K15198, JP23H04216, and JP24K09641 to M.T.; JP25K18695 to H. Kawai; JP21K19253 to H. Kawai and Y.H.; JP23K06310 to K.Y.; JP23K20044 and JP25K02371 to H.H.; JP26KJ0694 to Y.N.; JP25KJ1038 to K.S.; JP22H05094 and JP24K03266 to H.M.), JST BOOST Program Japan Grant Number JPMJBY25B1 to H. Kawai, Chan Zuckerberg Initiative napari Ecosystem Grants to H. Kawai, Japan Science and Technology Agency CREST (JPMJCR24T6 to Y.H.; JPMJCR21E2 to Y.H.), Japan Science and Technology Agency SPRING (JPMJSP2108 to Y.N.), the Japan Agency for Medical Research and Development (AMED) (JP19dm0207082 to Y.H. and JP21wm0525015 to H.M. and Y.H.; 256f0137010 to K.Y.; JP23wm0625001 and JP24wm0625103 to H.H.), Moonshot R&D from the Japan Science and Technology Agency (JST) (JPMJMS2024 to H.H.), SECOM Science and Technology Foundation Research grant (Y.H.), and The Uehara Memorial Foundation research grant (Y.H.) and Fusion Oriented Research for disruptive Science and Technology (FOREST) from JST (JPMJFR204D to H.H.) and Cooperative Study Programs of National Institute for Physiological Sciences (to Y.H.).

## Author contributions

Conceptualization: Y.H.

Methodology: Y.N., M.T., H. Kawai, K.S., K.Y., S.S., T.M., K.O., H.H., H.M.

Investigation: Y.N., M.T., K.L., H. Komaki, T.E., S.S., N.O., S.L., H.M., Y.H.

Visualization: Y.N., M.T., H. Kawai

Funding acquisition: H.K., H.H., N.O., H.M., Y.H. Project administration: Y.H.

Supervision: Y.H.

Writing – original draft: Y.N., M.T., H. Kawai, Y.H.

Writing – review & editing:

## Methods

### Mice

All animals were handled according to protocols approved by the Animal Care and Use Committee of The University of Tokyo (KA20-1) and Kumamoto University (A2026-033). Timed-pregnant ICR mice (Slc:ICR) were purchased from Japan SLC, Inc. (Shizuoka, Japan).

### Plasmids

The pCAG-mScarlet plasmid was generated by PCR amplification of the mScarlet coding sequence from pCAG-4xmt-mScarlet-iATPSnFR1.0^60^ using a forward primer (5′-CAAAGAATTGCTCGAGGCCACCATGAGCAAGGGCGAGGCAGTGAT-3′) and a reverse primer (5′-GAGTGAATTGCGGCCGCTTACTTGTACAGCTCGTCCATGC-3′).

The amplified fragment was inserted by In-Fusion cloning (Takara Bio) into pCAG-vGlut1-mCherry, a gift from Franck Polleux^61^, after linearization of the vector with XhoI and NotI-HF (New England Biolabs).

A plasmid encoding PPA-oxGFP-Halo-GPI (mPrion) was generated using PCR-based assembly. The coding sequence for the preproacrosin signal peptide (nucleotides 17–88 of NM_013455.4), oxGFP^37^ (Addgene #68069), HaloTag (Addgene #87645), and the GPI (mPrion) sequence (nucleotides 2401–2490 of NM_001278256.1, kindly provided by Drs. Miyawaki and Niino) were amplified and assembled into a single fusion construct. This cassette was inserted into pAAV2 SynTetOff-GFP^62^ by replacing the original GFP sequence, resulting in pAAV2 SynTetOff_PPA-oxGFP-Halo-GPI (mPrion). To generate pCAG-PPA-oxGFP-Halo-GPI (mPrion) with enhanced expression, the PPA-oxGFP-Halo-GPI (mPrion) sequence was amplified by PCR using PrimeSTAR Max DNA Polymerase (Takara Bio) with a forward primer (5′-AAAACTCGAGGCCACCATGGTAGAGATGCTG-3′) and a reverse primer (5′-AAAAGCGGCCGCTCATCCCACGATCAGGAAGATG-3′). The PCR product was cloned into the pCAG backbone by replacing the mScarlet sequence in pCAG-mScarlet plasmid, which had been digested with XhoI and NotI (New England Biolabs), using Ligation High Ver.2 (Toyobo).

The pTRE-Cre and pCAG-loxP-STOP-loxP-tdTomato-IRES-tTA-WPRE plasmids were constructed as previously described^44,63^.

### In utero electroporation

Pregnant mice at E15.5 were anesthetized with either a combination of medetomidine (0.3 mg/kg), midazolam (4 mg/kg), and butorphanol (5 mg/kg) or 2% isoflurane. Following laparotomy, the uterine horns were gently exteriorized. For EM analysis, a mixture of the vectors pCAG-PPA-oxGFP-HaloTag-GPI (1 µg/µl) and pCAG-mScarlet (0.5 µg/µl) and 0.01% Fast Green (Sigma) was injected into one side of ventricle of E15.5 embryos using a microinjector (BEX, BJ-120). For in vivo two-photon imaging, a plasmid solution containing pCAG-loxP-STOP-loxP-tdTomato-IRES-tTA-WPRE (1 μg/µl) and pTRE-Cre (5 ng/µl) was injected into the left lateral ventricle. Electroporation was performed to target cortical progenitors. Four 40-V pulses, each lasting 50 ms, were delivered at 950-ms intervals using Tweezer-type platinum electrode (diameter 5 mm; CUY650P5, Nepa Gene) connected to the GEB15 pulse generator (BEX). Following electroporation, the uterine horns were returned to the abdominal cavity, and the abdominal wall and skin were sutured. Atipamezole (Antisedan; Antisedan; 0.75 mg/kg) was administered to reverse medetomidine-induced anesthesia. The mice were then allowed to recover on a warming pad maintained at 37 °C.

### Sample fixation and sectioning

Following anesthesia, mice were heart perfused with 5 ml of the fixative solution containing 2% paraformaldehyde (EM grade, EMS) and 1% of Glutaraldehyde (GA, EMS) diluted in 0.1M Phosphate Buffer (PB). Brains were then dissected out and post-fixed in the same fixative solution for 2 hours. After embedded in low melting agarose (3%, MP Biomedicals), brains were sectioned at 100 µm thickness with a Leica VT1000S vibratome. For P2 samples, sections around the midline were selected, whereas for P5 samples, sections of the cerebral hemisphere contralateral to the electroporated hemisphere were selected.

### DAB staining

For HaloTag labeling, sections were incubated overnight at 4°C with a 1:500 dilution of HRP-conjugated HaloTag ligand in Phosphate Buffer Saline (PBS). The HRP-conjugated ligand was prepared using HaloTag Amine (O4) Ligand (P6741, Promega) and a Peroxidase Labeling Kit-NH2 (LK11, Dojindo) according to the manufacturer’s instructions and stored in 50% glycerol at –20°C until use. On the following day, the signal was amplified using a biotinylated tyramine (BT)-glucose oxidase (GO) reaction^38^. Sections were rinsed with 0.1 M PB and pre-incubated for 5 min at room temperature in 0.1 M PB containing 3 µg/ml GO (16831-14, Nacalai Tesque) and a 1:50,000 dilution of BT, protected from light. The reaction was initiated by the addition of β-D-glucose (G0047, TCI) to a final concentration of 2 mg/ml for 30 min. BT was synthesized by reacting 3.5 mg of Biotin-NHS (203112, Calbiochem) with 5 mg of tyramine hydrochloride (T2879, Sigma-Aldrich) in DMSO; the mixture was rotated overnight at room temperature, followed by the addition of 7.3 µl of monoethanolamine and further rotation for 4–5 h. Sections were then rinsed three times with PBS and incubated overnight at 4°C in an ABC solution. The ABC solution was prepared by mixing Reagent A and Reagent B (1:50 each; PK-6100, VECTASTAIN Elite ABC Standard Kit, Vector Laboratories) in PBS 30 min prior to use.

Following three washes in PBS, samples were incubated in 0.3 mg/mL DAB (049-22831, Wako) for 30 min, then supplemented with H_2_O_2_ (0.003% final concentration; 086-07445, Wako) for an additional 30 min, protected from light. After two PBS washes, samples were post-fixed with 1% GA in PBS at 4°C overnight.

### Preparation for electron microscopy imaging

To process for electron microscopy, tissue samples were osmicated using the reduced Osmium-Thiocarbohydrazide-Osmium (rOTO) staining method.

After washing with 0.1 M PB, samples were post-fixed with 1% Osmium tetroxide (OsO_4_) (Electron Microscopy Sciences), 1.5% potassium ferrocyanide (Fujifilm Wako pure chemical corporation, catalog No. 161-03742) in a 0.1 M PB for 90 min. After being rinsed 3 times with H_2_O, samples were stained with 1% thiocarbohydrazide (TCH) (Sigma-Aldrich) for 30 min. After being rinsed with H_2_O for three times, samples were stained with 1% OsO_4_ in H_2_O for 90 min. After being rinsed with H_2_O for two times at room temperature and three times with H_2_O at 50 °C, samples were treated with Walton’s lead aspartate (0.635% lead nitrate (Sigma-Aldrich), 0.4% aspartic acid (pH 5.2, Sigma-Aldrich)) at 50 °C for 20 min.

The sections were followed by incubations in an ascending ethanol series (5 minutes each in 50%, 70%, 80%, 90% ethanol/H_2_O), followed by three changes of 100% ethanol (5 min each), two changes of acetone (5 min each; Wako) and two changes of 5 min in n-butyl glycidyl ether (QY-1, Kanto Chemical). Samples were then infiltrated with a 1:1 mixture of QY-1 and Plain resin (Nissin EM) for 1 h at room temperature, followed by incubation in pure Plain resin at 4°C overnight. The following day, the samples were incubated in fresh resin for 2 h at room temperature. The sections were then flat-embedded between two sheets of Aclar film and polymerized at 70°C for 4 days.

### Immunohistochemistry

After in vivo two-photon imaging, brains were fixed with 4% paraformaldehyde in PBS, sequentially cryoprotected in 10%, 20%, and 30% sucrose, and cut into 20-μm-thick cryosections. Sections were blocked with 2% goat serum in PBS containing 0.1% Triton X-100 (PBS-T) for 1 h and incubated overnight at 4°C with rabbit anti-RFP antibody (1:1,000; PM005, MBL). After three 10-min washes with PBS-T, sections were incubated with Alexa Fluor 594–conjugated donkey anti-rabbit IgG (1:1,000; A21207, Invitrogen), and nuclei were counterstained with Hoechst 33342 (080-09981, Fujifilm Wako). Sections were washed three times with PBS-T and mounted. The sections were examined post hoc to assess the aspirated region and tdTomato-positive axons in the white matter.

### Confocal microscopy

For analysis of HaloTag localization in EM samples, brain slices were labeled with HaloTag SaraFluor 650T Ligand (GORYO Chemical) and imaged using a CFI Plan Apochromat Lambda D 100× oil-immersion objective (NA 1.45). For post hoc analysis of tdTomato-positive growing axons in the white matter following in vivo two-photon imaging, immunostained sections were imaged using a CFI Plan Apochromat Lambda 20× objective (NA 0.75) or a CFI Apochromat LWD 40× WI Lambda S objective (NA 1.15). Images were acquired on a Nikon Ti2 Eclipse microscope with an A1R confocal, a laser unit (Nikon; LU-N4, 405, 488, 561, and 640 nm), and filters (450/50 nm, 525/50 nm, 595/50 nm, 700/75 nm for 405 nm, 488 nm, 561 nm, 640 nm laser, respectively). All equipment was controlled via NIS Elements software (Nikon). The resulting images were deconvoluted with NIS-elements (Nikon) and processed with NIS-elements (Nikon) or ImageJ.

### Electron microscopy

Sample blocks were trimmed with a TrimTool diamond knife (Trim 45; DiATOME) and ultrathin sectioned with an Ultra Diamond Knife (Ultra 45°; DiATOME) in a Leica Ultramicrotome (UC7). Sections were collected on the cleaned silicon wafer strip and imaged with a scanning electron microscope (JSM-IT800SHL; JEOL), using the Scintillator Backscattered Electron Detector. Images were acquired at a resolution of 5 nm/pixel.

For volumetric analysis, serial block-face scanning electron microscopy (SBF-SEM) was performed using either a Merlin field-emission scanning electron microscope (Carl Zeiss) equipped with a 3View in-chamber ultramicrotome and backscattered electron detector (Gatan, Pleasanton, CA, USA), or a Teneo VS field-emission scanning electron microscope (FEI, Thermo Fisher Scientific) equipped with a VolumeScope in-chamber ultramicrotome.

For imaging with the Merlin–3View system, sample blocks were mounted on aluminum rivets using conductive TK Paste (Kaken Tech) and gold-sputter-coated to increase conductivity. Serial images were acquired at an accelerating voltage of 1.3-1.4 kV and a dwell time of 1 µs per pixel with a pixel size of 5 nm with a section thickness of 25–30 nm.

For imaging with the Teneo VS–VolumeScope system, resin blocks were trimmed, mounted onto aluminum SBF-SEM specimen stubs using conductive epoxy resin, and platinum-coated to minimize charging artifacts. Serial images were acquired using a T1 backscattered electron detector at an accelerating voltage of 1.73 kV and a dwell time of 1 µs per pixel. The section thickness was 40 nm, and the lateral pixel size was 5 or 7 nm.

Serial electron micrographs were aligned using the Linear Stack Alignment with Scale-Invariant Feature Transform (SIFT) plugin implemented in ImageJ.

### Segmentation of vEM datasets

Axons were traced with the original Segment Anything Model (SAM^29^; ViT-H checkpoint) used zero-shot with its weights frozen, that is, without fine-tuning or any other update to its parameters. No adapter or auxiliary network was trained, and no task-specific training data were used. After one bounding box on a seed section was drawn, every prompt is generated by the pipeline from that section onward. On each section SAM was run on a 1,024 × 1,024-pixel region of interest centered on the incoming box and queried with multiple mask outputs; among the returned candidates, the mask with the highest intersection over union (IoU) against the preceding section’s mask was retained. Prompting was by bounding box alone. If that mask had both recall below 0.75 and precision below 0.6 against the preceding mask, indicating a jump onto a different profile rather than a taper (which lowers recall but not precision), SAM was re-run with the preceding mask as a low-resolution logit prior, and the better-matching of the two results was kept. If its recall remained below 0.40, the profile was deemed to have ended and the trace was terminated.

In the refined condition, the retained mask was then improved by re-prompting SAM from its own output within the same section: at each iteration the mask was split into connected components, each was re-segmented from its own tight bounding box, and the results were combined, stopping when the IoU between successive masks exceeded 0.99 or after ten iterations. Iterations that retained at least 80 % of the one-pass mask and did not exceed three times its area were trimmed at dark membrane boundaries, using a Sato ridge filter (scikit-image 0.26; sigmas 1–3, black_ridges = True) to remove in-mask pixels above the 70th percentile of the ridge response, with trimming skipped below 200 pixels. The largest trimmed mask was accepted, or the one-pass mask itself if none exceeded it. Non-largest connected components were then discarded when they exceeded 500 pixels, smaller ones being retained. The tight bounding box of the resulting mask prompted the next section, whereas only its largest connected component was written to the output stack. The two benchmark conditions differ only in this refinement step and its membrane gate; all other processing is identical.

To benchmark automated tracing, 50 unlabeled axons were selected at random from a 100-section SBF-SEM volume. Each segmentation trial was initialized with one expert-drawn bounding box on the first section and traced automatically to the final section. Every axon was traced under both conditions from the same initial box, with the outer loop or CLAMP. Because the method propagates a single connected profile, axons that branched into two or more profiles within the traced range were excluded and replaced by further random draws. Ground truth was drawn by an expert as a bounding box around each target profile on the final section, without reference to the automated output. The tight box of the automated final-section mask was compared with the ground truth by IoU; IoU ≥ 0.7 defined success and an empty mask scored zero. Success was also tabulated from 0.3 to 0.9 in steps of 0.05 (Fig. 2d). Because ground truth exists only on the final section, this is an endpoint success proxy and does not validate intermediate sections. The benchmark axons carried no DAB label, so tracing relied on membrane contrast alone rather than on the electron-dense staining available in the labeled reconstructions. Tracing was run in PyTorch 2.10 with Metal Performance Shaders (MPS) acceleration on an Apple M5 MacBook Pro (24 GB unified memory, macOS 26).

To assess robustness to prompt placement (**Extended Data Fig. 1**), 24 sections were sampled from axons that could be followed continuously through the volume. For each section, the bounding-box prompt was displaced in four directions by the indicated fraction of the box side length. Segmentation was performed either with a single SAM inference (– inner loop) or with the inner refinement loop run to convergence within the same section (+ inner loop). This analysis isolates the box-refinement loop itself: SAM returned a single mask, and no coverage guard, membrane gate or component filtering was applied. The resulting masks were compared by IoU with the reference mask generated from the undisplaced prompt. Values obtained from segmentations using prompts displaced in four directions were averaged to yield a single value per section. The undisplaced prompt was defined as a tight bounding box around the converged mask. A single SAM call using this prompt reproduced the same mask exactly, therefore, both conditions scored 1.0 at zero displacement. Because this reference is the pipeline’s own output rather than a manual segmentation, the analysis measures robustness to prompt placement, not absolute accuracy. EM traces were inspected against the raw images; those in which propagation failed, whether the mask was lost, collapsed, or transferred to a neighboring profile, were corrected by re-seeding from the last correct section or by editing the affected masks in napari. Corrected traces were used for the morphometric analyses. The benchmark in Fig. 2d used uncorrected output.

### Tracking and segmentation of time-lapse data

For in vivo time-lapse data the per-frame prompt came from VitTracker (cv::TrackerVit in OpenCV 4.13, object_tracking_vittrack model from the OpenCV Model Zoo, https://github.com/opencv/opencv_zoo) initialized from one user-drawn box on the first frame and passed to the same SAM. Refinement was not applied in this mode, and all masks were manually proofread, with corrections made by re-seeding from the last correct frame or by editing the masks in napari. Growth cones were sufficiently sparse in the two-photon imaging fields that SAM did not inadvertently segment neighboring growth cones. Instead, tracking was limited by inter-frame movement and shape change.

### Morphometric analysis of reconstructed growth cones

Filopodia were identified from 3D segmentations as axon-connected protrusions with a skeleton path length greater than 1 µm and a maximum Euclidean-distance-transform-based thickness below 300 nm. Filopodia contacting the axon shaft within 5 µm of the axon tip were counted. The analysis included P2 growth cones, n = 27 structures from N = 3 mice (2, 1, and 24 growth cones per mouse), and P5 growth cones, n = 11 structures from N = 1 mouse. One growth cone was treated as one statistical observation, and filopodia numbers were compared between P2 and P5 using a two-sided Mann–Whitney U test.

Axonal center lines were reconstructed from 3D annotations and used to sample perpendicular cross-sections at 100 nm intervals. Growth cones were analyzed over the distal 5 µm, whereas control shafts were analyzed over their annotated lengths (> 5 µm) after excluding 100 nm from each end. Equivalent circular diameter was calculated from cross-sectional area. Cross-sectional aspect ratio was calculated as the square root of the ratio between the major and minor eigenvalues of the area-weighted 2D covariance matrix. The mean equivalent diameter and mean aspect ratio across valid sections were used as the representative values for each structure. The analysis comprised P2 growth cones, n = 27 structures from N = 3 mice; P5 growth cones, n = 11 structures from N = 1 mouse; P2 control shafts, n = 10 structures from N = 1 mouse; and P5 control shafts, n = 6 structures from N = 1 mouse. Equivalent diameters of the four groups were compared using Welch’s one-way ANOVA followed by Games–Howell pairwise comparisons. Aspect ratios of control shafts and growth cones were compared separately at P2 and P5 using two-sided Welch’s unpaired t-tests.

Orientation analyses included filopodia meeting the morphological criteria above and contacting the axon shaft within 10 µm of the axon tip. This broader window retained growth cone-associated filopodia with attachment sites slightly proximal to the distal 5 µm growth cone region.

The analysis included n = 35 P5 filopodia nested within 9 growth cones from one EM dataset and N = 1 mouse. For each growth cone, the growth cone plane was estimated by 3D principal-component analysis of surface voxels within the distal 5 µm. The first two principal components defined the growth cone plane, and PC3 represented its normal. Filopodium direction was represented by the 3D chord from its shaft-contact end to its distal endpoint. The unsigned angle between the filopodium and the growth cone plane ranged from 0°, indicatin an in-plane orientation, to 90°, indicating alignment with the plane normal. Planar alignment was quantified from the cosine of twice each plane angle. Statistical significance was evaluated using a one-sided Monte Carlo test against isotropically distributed 3D orientations (10^6^ replicates). This 3D null was used because uniformly distributed angles between 0° and 90° do not represent isotropic orientations in three-dimensional space.

The projected orientation of each growth cone plane was represented by its surface-PCA PC2 axis projected onto the imaging XY plane. Angles were registered to the anatomical dorsoventral (DV) axis using dataset-specific offsets determined from landmarks in low-resolution electron micrographs. Bias toward the DV axis was assessed using one-sided axial V-tests toward 0°, with equal weights and unadjusted p values.

Each 3D filopodium chord was projected onto the imaging XY plane and registered to the DV axis. Bias toward the DV axis was assessed using a one-sided axial V-test toward 0°. Axial angles were doubled before testing, observations were assigned equal weights, and the reported p value was unadjusted.

### In vivo time-lapse imaging by using 2-photon microscopy

At postnatal day 4 (P4), tdTomato fluorescence was examined using either an upright fluorescence microscope (Leica) or a Twin LED Light (Relyon), and pups exhibiting detectable fluorescence were selected for cranial-window surgery. Under isoflurane anesthesia, a small cranial window approximately 500 μm in diameter was created, and a custom-made head bar was attached to the adjacent skull. After removal of the skull and dura mater, the overlying cortical tissue was gently aspirated using an aspirator to a depth of approximately 500–600 μm from the brain surface. Bleeding from the exposed cortex was controlled using Spongel (Astellas Pharma). After bleeding from the resected area had ceased, the exposed tissue was covered with 1% agarose dissolved in artificial cerebrospinal fluid (ACSF), and the imaging window was sealed with a glass coverslip. The coverslip was secured in place with cyanoacrylate adhesive (Aron Alpha), and its edges were sealed with dental cement.

Pups were allowed to recover from anesthesia on a heating pad for at least 1 h. No overt signs of postoperative distress were observed; the pups suckled normally and behaved similarly to untreated littermates. Before imaging session, a custom-made 3D-printed imaging cup was attached and sealed with dental cement to retain the immersion water. During imaging, mice were anesthetized with 0.7%–0.9% isoflurane. In vivo images were acquired using a two-photon microscope (Leica SP8MP or A1R, Nikon) equipped with an ALCOR-920 fibre laser (Spark lasers) or a Mai Tai eHP DeepSee Ti laser (Spectra-Physics) tuned to 920 nm, a GaAsP detector, and a 25× objective lens (NA 0.95 or 1.1). Between imaging sessions, pups were housed with their littermates in a small cage placed on a heating pad maintained at 37 °C. All pups remained healthy throughout the imaging experiments.

### Registration of in vivo two-photon time-lapse images (related to *Figure 4f,g*)

To correct for three-dimensional motion between time points, the acquired image stacks were registered before quantitative analysis. At each time point, the best-focused optical sections were manually selected from a z-stack of approximately 40 sections, yielding a total of approximately 800 two-dimensional images from eight axons for quantitative analysis.

Estimation and correction of lateral displacements of the two-dimensional time series was performed with a custom Python pipeline (Python 3.14.0) built on the SOFIMA library^64^. Lateral displacements between consecutive frames were first estimated by masked normalized cross-correlation of local image patches (patch size = 80 pixels). To capture both large and small displacements, estimation proceeded coarse-to-fine over eight passes on a Gaussian-antialiased image pyramid (full, 2-fold and 4-fold downsampled; scikit-image 0.26.0), with the spacing between patch centers decreasing from 640 to 20 pixels. In each pass the displacement field obtained in the preceding pass was used to pre-target the correlation search, and the two fields were then reconciled after resampling the coarser field onto the finer grid. Flow vectors were rejected when the sharpness of the correlation peak, or its ratio to the next-highest peak, fell below 1.6, or when a vector deviated by more than 20 pixels from the local median. The frame-to-frame displacement was then taken as the median of all accepted flow vectors, and these displacements were summed over the time series to give the shift of each frame relative to the first time point. To avoid artificial rotation or unstable warping, only the translation was corrected, and no rotational or non-rigid component was estimated. Shifts were applied as integer-pixel translations without interpolation, so that pixel values were left unchanged; regions translated out of the field of view were left empty and excluded from analysis.

### Visualization of growth cone centroid trajectories (related to *Figure 4h*)

Growth cone centroid coordinates were obtained from the proofread binary masks at each time point. Coordinates were expressed relative to the centroid position in the first frame, and consecutive positions were connected in temporal order. Individual positions were color-coded according to elapsed imaging time, with the first and last positions indicated separately. Trajectories were displayed without rotational alignment, and the y-axis was inverted to preserve the orientation of the original image. Coordinates are expressed in µm.

### Quantification of growth cone velocity (related to *Figure 4i*, Extended Data Figure 5)

For each axon, a fixed global progression axis was defined as the direction of displacement between the growth cone centroids in the first and last analyzed frames and was held constant throughout the image sequence. The coordinates of all pixels within each proofread growth cone mask were projected onto this axis. The anterior position of the growth cone was defined as the 90th percentile of the projected mask-pixel positions (P90), such that 90% of the projected pixels were located posterior to this position and 10% anterior to it (**Extended Data Figure 5**). Growth cone displacement between consecutive frames was calculated as the difference in P90 position, and velocity was calculated by dividing the displacement by the imaging interval (Δt = 1.5 min). Positions and velocities are reported in µm and µm/min, respectively.

### Visualization and classification of growth cone movement states (related to *Figure 4i*, j)

Each valid frame-to-frame interval was classified into one of three movement states according to its velocity: Advancing: v > 0.30 µm/min, Pausing: |v| ≤ 0.30 µm/min, Retracting: v < −0.30 µm/min. For the representative trace in Figure 4i, analysis was restricted to the first uninterrupted sequence of valid velocity intervals and terminated at the first subsequent missing or excluded interval. To visualize the longer-term velocity trend, a centred five-frame moving average was calculated. The movement-state classification was based on the raw frame-to-frame velocity. Movement-state periods were displayed as a color-coded band above the velocity trace, with advancing, pausing, and retracting shown in magenta, gray, and blue, respectively. For the population-level analyses in Figure 4j, all valid intervals from each axon were included. Consecutive intervals assigned to the same state were treated as a single bout. Missing or excluded velocities and gaps in frame numbering terminated a bout. Bout duration was calculated as the number of consecutive intervals multiplied by 1.5 min. For each axon, the proportion of valid observation intervals assigned to each state was calculated separately, and group values are reported as the mean ± SEM across axons.

### Quantification of growth cone morphological features (related to *Figures 5a-d*, Extended Data Figures 6 and 7)

Circularity, eccentricity, and area were calculated from the proofread binary growth cone masks. Area was calculated as the number of pixels within each mask multiplied by the calibrated pixel area. Circularity was calculated as:

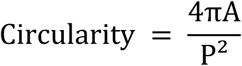

where A is the mask area and P is its perimeter.

Eccentricity was calculated from an ellipse fitted to the second central moments of the mask:

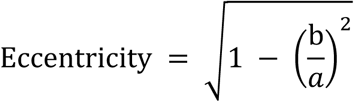

where a and b are the semi-major and semi-minor axes of the fitted ellipse, respectively.

### Correlation analyses of growth cone morphology with velocity (related to *Figure 5b* and Extended Data Figures 6 and 7)

Target time points were included when a complete sequence of four consecutive frames from *t*_−1_ to *t*_2_ was available within the same kinematic segment. No event-selection criterion based on velocity magnitude or direction was applied. Associations of circularity, eccentricity, and area at *t_−1_*, *t_0_*, and *t_1_* with velocity were evaluated using Pearson’s correlation coefficients. Corresponding 95% confidence intervals were calculated for each coefficient, and ordinary least-squares regression lines were displayed for visualization.

### Analysis of circularity transitions and subsequent velocity (related to *Figure 5c–g*)

Events were included when consecutive frames spanning *t_0_* to *t_2_* were available within the same kinematic segment. Circularity quartiles at *t_0_* and *t_1_* were calculated separately across all eligible events. Events were classified as Cir^Low→Low^, Cir^Low→High^, Cir^High→Low^ or Cir^High→High^ when circularity fell within Q_1_ or Q_4_ at both time points. Transition histories were compared separately among events reaching the same Cir(*t_1_*) endpoint: Cir^Low→High^ versus Cir^High→High^ for the Q_4_ endpoint and Cir^Low→Low^ versus Cir^High→Low^ for the Q_1_ endpoint. Within-group changes from *v_0_* to *v_1_* were evaluated using paired *t*-tests. Between-group differences in *v_1_* were evaluated using linear mixed-effects models fitted by maximum likelihood, with transition group as a fixed effect and axon identity as a random intercept:

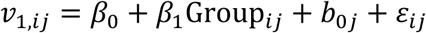

where *b_0j_* denotes the random intercept for axon *j*. The transition-group effect was tested using a one-degree-of-freedom likelihood-ratio test against a reduced model without the group term:

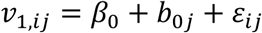

To determine whether the between-group difference in *v_1_* persisted after adjustment for *v_0_*, an additional model included *v_0_* as a fixed effect:

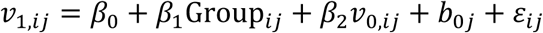

The transition-group effect was evaluated using a one-degree-of-freedom likelihood-ratio test against the reduced model:

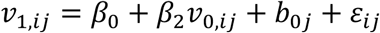

### Comparison of circularity endpoints within each initial state (related to Extended Data Figure 8)

The event-selection criteria and circularity-quartile definitions were identical to those used for the circularity-transition analysis in **Figure 5**. The four transition groups were pooled according to their Cir(*t_1_*) endpoint, irrespective of Cir(*t_0_*): the Cir(*t_1_*)^Low^ group comprised Cir^Low→Low^ and Cir^High→Low^ events, whereas the Cir(*t_1_*)^High^ group comprised Cir^Low→High^ and Cir^High→High^ events. Differences in *v_1_* between the two endpoint groups were evaluated using a linear mixed-effects model fitted by maximum likelihood, with Cir(*t_1_*) endpoint as a fixed effect and models containing Cir(*t_1_*) endpoint and axon identity as a random intercept:

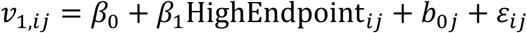

The Cir(*t_1_*) endpoint effect was evaluated using a one-degree-of-freedom likelihood-ratio test against the corresponding reduced model lacking the endpoint term:

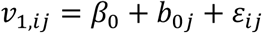

The coefficient for *HighEndpoint* represents the model-estimated difference in *v_1_* between events ending in Q_4_ and those ending in Q_1_, accounting for the clustering of events within axons.

### Statistics

Tracing conditions were compared on a paired, per-axon basis: paired success counts at a given IoU threshold with an exact McNemar test (a two-sided binomial test on the discordant pairs), and paired per-axon IoU values with a two-sided Wilcoxon signed-rank test. No correction for multiple thresholds was applied; IoU ≥ 0.7 was designated the primary criterion in advance.

## Data availability

SBF-SEM data are available on EMPIAR under accession code EMPIAR-14003 (https://doi.org/10.6019/EMPIAR-14003).

In vivo two-photon time-lapse recordings are available from the corresponding author on reasonable request. Source Data are provided with this paper for the measurements underlying Figs 2d, 3e–l and 5b–g and Extended Data Figs 1 and 6–12.

## Code availability

CLAMP and the VitTracker-guided prompt generation used here are implemented in an open-source napari plugin released under the Apache-2.0 licence. The plugin and the analysis code that reproduces the figures will be made publicly available on GitHub and archived with a DOI at Zenodo before publication, and are available to editors and reviewers on request during review.

